# PPIX-binding Proteins Reveal Porphyrin Synthesis and Ferroptosis Link

**DOI:** 10.1101/2022.06.01.494336

**Authors:** John Lynch, Yao Wang, Yuxin Li, Kanisha Kavdia, Yu Fukuda, Sabina Ranjit, Camenzind G. Robinson, Christy R. Grace, Youlin Xia, Junmin Peng, John D. Schuetz

## Abstract

All aerobic organisms require the cofactor heme to survive, but its synthesis requires formation of a potentially toxic intermediate protoporphyrin IX (PPIX). Little is known about the extent of PPIX’s cellular interactions. Here, we report the development of PPB, a biotin-conjugated, PPIX-probe that captures proteins capable of interacting with PPIX. Quantitative proteomics with PPB identified common proteins among a diverse panel of mammalian cell lineages. Pathway and quantitative difference analysis revealed PPB-bound proteins related to iron and heme metabolism and suggested that these processes might be altered by heme/porphyrin synthesis. We show that increased heme/porphyrin synthesis in cells promotes ferroptosis that is pharmacologically distinct from canonical ferroptosis driven by erastin, an inhibitor of the cystine/glutamate antiporter. Proteomic data derived from PPB revealed an interactor, PRDX3, a mitochondrial peroxidase, that modulated heme/porphyrin biosynthesis driven ferroptosis. Consistent with a role in porphyrin-induced ferroptotic death targeted gene knockdown of PRDX3, but not peroxidases, PRDX1 or 2, enhanced porphyrin-induced ferroptotic death. The relationship between increased heme/porphyrin synthesis and ferroptosis was also found in a ferrochelatase-deficient T-lymphoblastoid leukemia cell line, suggesting potential strategy for treating certain cancers. We demonstrate that when the PPB probe is coupled with unbiased proteomics a previously unreported relationship between heme/porphyrin synthesis, and ferroptosis was discovered.

## Introduction

Heme, a prosthetic group for enzymes in key cellular processes, such as electron transport, detoxification, protection against oxygen radicals, and oxygen transport^1–3, 4,5^ Additionally, heme can bind transcription factors to regulate their activity^6, 7^. Heme synthesis requires protoporphyrin IX (PPIX), the potentially toxic penultimate product^8^. PPIX may accumulate in cells for a variety of reasons, including impairment of the terminal pathway enzyme, ferrochelatase (FECH), which inserts ferrous iron into PPIX to form heme^9–11^. Cancer cells show enhanced levels of PPIX, often due to reduced levels of FECH^11–13^. PPIX and heme formation are promoted following exogenous addition of a precursor in the heme/porphyrin biosynthetic pathway, delta-aminolevulinic acid (ALA)^12, 14^. This property has been exploited to kill cancer cells, with the elevated PPIX levels promoting cell death^15^, in combination with light. This phototoxicity induced by PPIX has long been recognized and has been highly studied as a source of toxicity. Elevated PPIX levels, as is the case in porphyrias^16^, a disease where porphyrin precursors, especially PPIX accumulate as a result of either genetic deficiencies of the synthetic enzymes in the heme pathway^17, 18^. In the absence of phototoxicity, the determination of PPIX-interacting proteins might uncover how PPIX elicits cell death and affects proliferation. Such knowledge will be beneficial to developing strategies to enhance cancer therapies, or to ameliorate suffering from porphyria^19, 20^, or from the elevated porphyrin concentrations that occurr secondary to environmental toxicants (e.g., lead, arsenic) and other chemicals that disrupt heme biosynthesis in mammals^21^. The mechanism for PPIX toxicity, in the absence of light^22^, is unclear^23^. Does PPIX cytotoxicity relate to reactive oxygen species (ROS), protein interactions, dysregulation of certain intracellular processes or a combination of both? Some data suggests that, PPIX cytotoxicity is independent of photoactivation (e.g., TP53)^24^. In macrophages, the voltage dependent anion channel (VDAC) PPIX interaction elicits mitochondrial permeability pore opening, a lethal event in some cell types^25^. Insights into the mechanisms of PPIX toxicity are hampered by the paucity of protein interaction knowledge.

PPIX is also intriguing from the perspective of intracellular trafficking. PPIX is formed in the mitochondrial matrix^26^, yet distributes throughout the cell to the plasma membrane, cytosol, mitochondria, endoplasmic reticulum, Golgi, and nuclear envelope^25, 27–33^, suggesting that it is either actively exported or chaperoned from the mitochondria after its synthesis in the mitochondrial matrix. Intracellular PPIX transit likely occurs by protein interactions (e.g., chaperones) that we hypothesize might be conserved among different cell lineages. For instance, it is known that PPIX binds to the mitochondrial peripheral benzodiazepine receptor^34^ and is exported from the cell by the plasma membrane ABC transporter, ABCG2^35–38^.

It is unclear as to how many cellular proteins PPIX interacts with, and whether interacting proteins are common among different cell lineages or if they vary in their potential for PPIX interaction. Additionally, the proteins involved in PPIX interactions might be conserved between species. Hence, identifying proteins that interact with PPIX might be the initial step in facilitating the discovery of new biological and functional relationships. Our approach to identify PPIX-binding proteins was to develop a probe conjugating biotin to PPIX (referred to as PPB); this unique conjugate was substantiated by its interactions with albumin as does PPIX^39^. The PPB bound proteins were readily captured and this approach minimized the non-specific interactions of proteins with PPB by employing biotin instead of PPIX, as competitor because of the latter’s potential to interact and trap proteins at high concentrations. This approach was then used to evaluate PPIX-binding proteins in multiple cell lineages, which were then quantitatively determined by a non-labeled approach of Tandem mass tag (TMT) proteomics^40^. We identified multiple PPB binding proteins that are distributed in multiple subcellular compartments. Importantly, these interacting proteins were quantitatively different but common across cell lineages employed and revealed homologous mouse and human proteins of, suggesting conservation among PPB-interacting proteins. Our data analysis revealed distinctions between the interacting proteins in NIH3T3 cells vs other cell lineages. The NIH3T3 cells displayed a higher accumulation of PPIX when heme/porphyrin synthesis was activated by ALA exposure. Additionally, we found that some PPB-interacting proteins were common to the heme biosynthetic and ferroptosis pathways. Furthermore, treatment of NIH3T3 cells with ALA produced toxicity through an iron-and ROS-mediated pathway consistent with ferroptosis. This observation led to the elucidation of a previously unknown relationship between heme/porphyrin biosynthesis and ferroptosis. We hypothesized that ROS produced during PPIX formation might be mitigated by antioxidant enzymes. Our screen revealed that genetic suppression of one such PPB binder, the enzyme PRDX3, increased the toxicity of ALA in NIH3T3. This PPIX-mediated ferroptotic pathway was predictively found to be active in Jurkat, a T-cell leukemia cell line with ferrochelatase deficiency, suggesting this knowledge might be of therapeutic importance.

Together our results show that by combining biotin with PPIX, a small molecular probe, PPB, can exploit proteomic techniques to identify interacting proteins in multiple cell types, and facilitate the discovery of a connection between porphyrin/heme biosynthesis and the ferroptosis cell death pathway. Further, this knowledge of the interacting proteins enabled the identification of an unknown modulator of this ferroptosis pathway, mitochondrial peroxidase, PRDX3, a gene that would not likely not have been otherwise identified. It is likely that a similar approach could be used to identify other hidden relationships between biologically active molecules and their metabolic pathways.

## Results

### Development of a probe to identify cellular Protoporphyrin IX (PPIX)-binding proteins

Despite the ubiquity of heme synthesis in mammalian cells, there is no comprehensive knowledge of the intracellular proteins that interact with the penultimate intermediate in its synthesis, PPIX. It is known that PPIX synthesis occurs in the mitochondrial matrix, from which it may be expelled from mitochondria by an unknown process, and it is expelled from cells by the transporter ABCG2 and possibly other transporters (**Fig. 1a**). To address this gap, we developed a probe, PPB, to identify PPIX-binding proteins. We reasoned that a biotin moiety joined to PPIX, but tethered by a 13-atom-long flexible linker including two polyethylene glycolsand attached by a carboxamide linkermight capture PPIX-binding proteins (**Fig. 1b**). The NOESY and TOCSY spectra with chemical shift assignments (**Supplemental Data Fig 1 and 2 and Table 1**) show that the hydrogens at N8 and N7 demarcate the PEG linker connection to biotin (hydrogens at 49 & 56) and to PPIX [hydrogen at 32 (methyl)] respectively. These analyses are consistent with the proposed structure (**Fig 1b**). Because of PPIX symmetry, this analysis is unable to distinguish if the probe is attached to either C13 or C17 position of PPIX propionate sidechain. An advantage of this long-linker design is that potential PPIX binding proteins might be less likely to be sterically restricted compared to previous probes attached directly to a bead such as was done with the commercial reagent for heme binding proteins, hemin-agarose.

**Fig 1.**
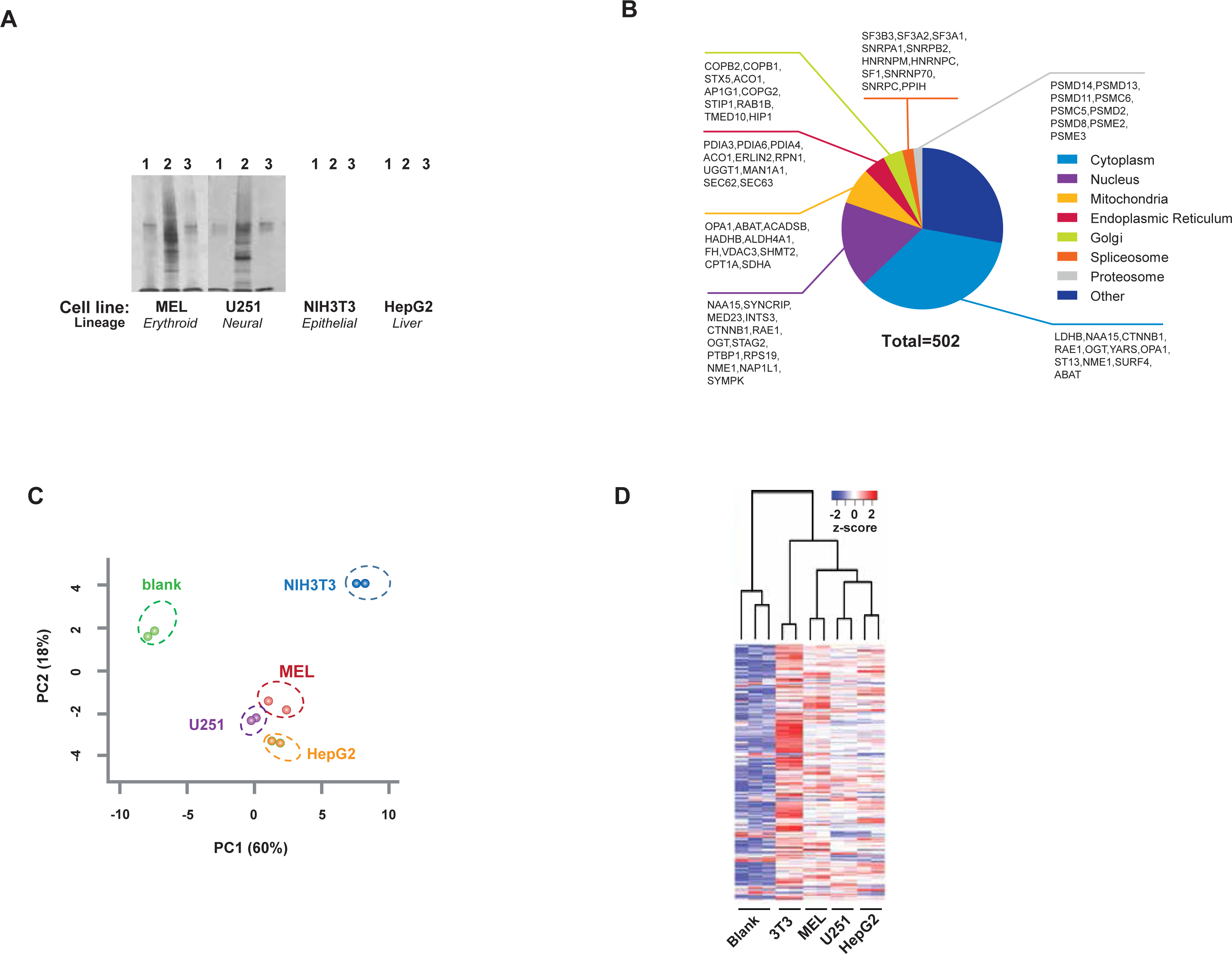
Mass spectrometry using tandem mass tag labeling indicates cellular differences in PPIX-binding proteins. **A,** Representative silver stain of PPB immunoprecipitation and controls from four cell lines. **B,** Principal component analysis (PCA) of TMT proteomics data. **C,** Unsupervised hierarchical clustering of cell lines’ quantitative TMT proteomic data from PPB immunoprecipitationassay.**D,** Heat map of PPB proteins from four different lineage cell lines.

**Fig 2.**
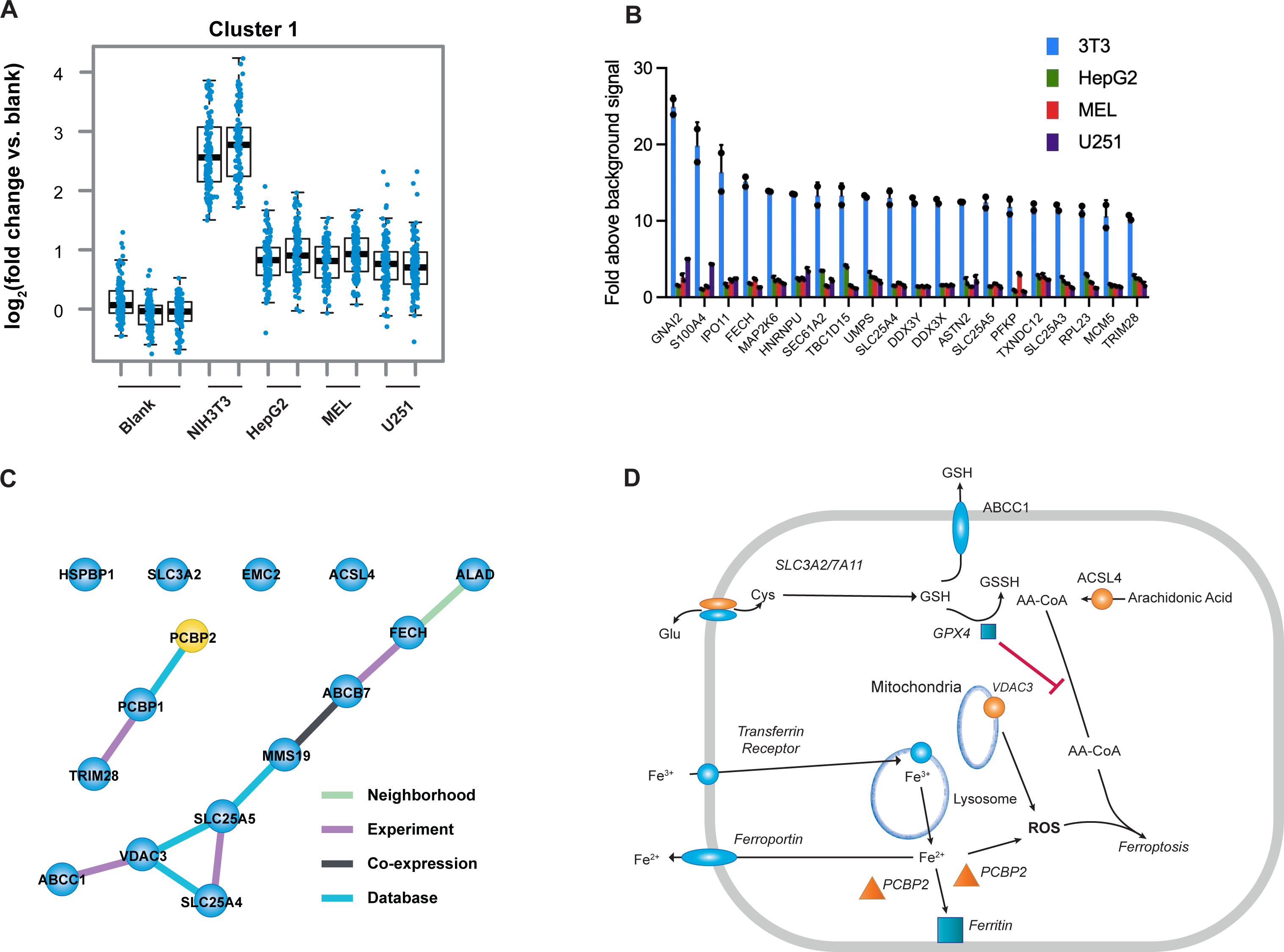
Analysis of PPIX-interacting proteins identifies NIH3T3-specific cluster of PPIX-binding proteins related to iron and mitochondrial metabolism. **A,** Weighted gene co-expression network analysis (WGCNA) of the PPB and tandem mass tag (TMT) quantitative proteomics data identified a cluster of proteins enriched in NIH3T3 cells. **B,**Highest candidate PPIX-binding proteins from NIH-3T3 cluster 1 shown relative to other cell lines (mean ± SEM, n=2 biologically independent samples)**. C,** STRING protein-protein interaction (PPI) network analysis of cluster 1 identifies a protein module enriched for iron and oxidative phosphorylation. **D,** PPIX-binding proteins identified, demonstrating a role in iron pathway and ferroptosis.

The molecular weight was determined by mass spectrometry and the structure of PPB was confirmed with these 2D NMR spectra: [^1^H, ^1^H]-TOCSY, [^1^H, ^1^H]-NOESY, [^1^H, ^1^H]-COSY, [^15^N, ^1^H]-HSQC and [^13^C, ^1^H]-HSQC (**Supplemental Data Fig 1 and 2 and Fig. 1c**). We anticipated that the lack of hindrance of PPIX and its physical separation from the biotin moiety would allow capture of PPIX-binding proteins by avidin beads (**Fig. 1d**). To reduce non-specific protein interactions, we tested 50-fold excess biotin to overwhelm the binding capacity of the avidin beads or excess free PPIX to block the capture of proteins as described below (**Supplemental Data Fig. 3**).

**Fig 3.**
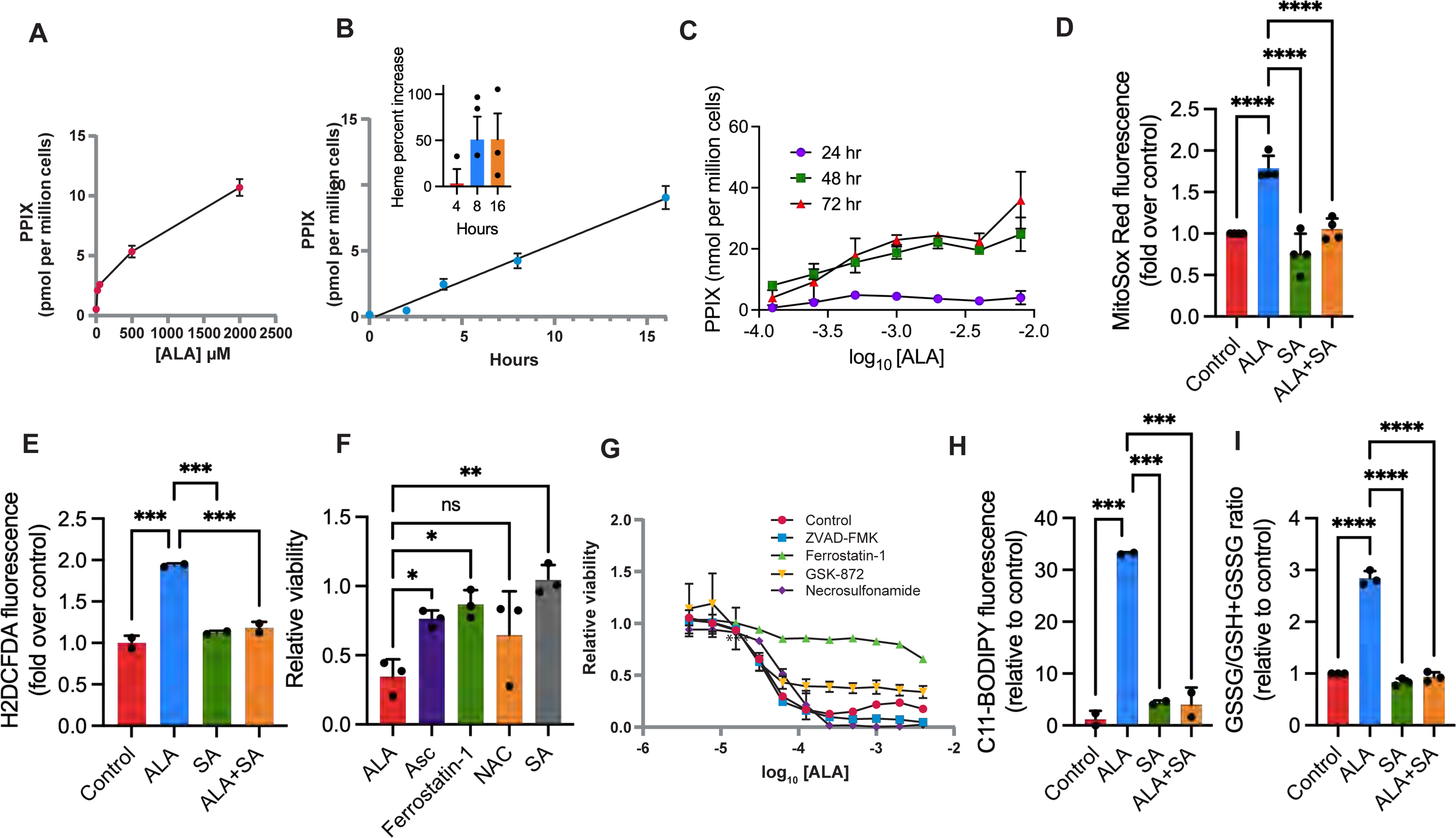
ALA induces increases ROS and promotes ferroptosis. **A,** ALA enhances generation of PPIX in NIH3T3 cells in a dose-dependent manner following treatment with various ALA concentrations. (n=2 independent samples**). B,** Dose-dependence of PPIX and heme (mean+/SEM) increase during ALA incubation, (n=2 samples, representative of 3 independent experiments for PPIX, n=3 independent experiments for heme). **C,** Release of PPIX into the media by ALA treated NIH3T3 cells at 24 (purple circles), 48 (green squares), and 72 (red triangles) hours. **D,** Mitochondrial superoxide levels (mean ± SEM) in ALA-treated cells, as measured by MitoSOX red fluorescence (n=4 independent experiments). **E,** Total cellular ROS in ALA-treated cells, as measured by DCFDA fluorescence (average of 3 independentexperiments). **F,** ALA cytotoxicity (mean ± SEM) is suppressed by antioxidants; ascorbic acid (500 µM), N-acetyl cysteine (5 mM) or Ferrostatin-1 (10 µM) (n=3 samples representative of at least 3 independent experiments forall drugs tested). **G,** Effect on ALA cytotoxicity (mean ± SEM) b y inhibitors of apoptosis, ferroptosis, and necroptosis (n=2 independent samples). **H,** Increased lipid peroxidation (mean ± SEM) with ALA treatment and attenuation by SA, as assessed by C11-Bodipy (n=2 samples, representative of 3 independent experiments).**I,** Oxidized glutathione (mean ± SEM) changes in ALA-treated cells (n=3 independent experiments). Significance is indicated by the number of attached asterisks. \**P* < 0.05, \*\**P* < 0.01, \*\*\**P* < 0.001, ****P<0.0001, and ns indicates no significance. Significance in Figure 3 were evaluated using one-way ANOVA, while paired Student’s *t*-test, two-sided, was used to determine significance between curves in Figure 3F.

### PPIX biotin probe (PPB) validation

To evaluate the PPIX binding activity of our probe PPB, we tested its interaction with albumin, a known PPIX-binding protein with a single binding site. To prevent photo-oxidation of either PPB or PPIX, incubations were conducted in the dark. Albumin’s Trp214 residue interacts with PPIX, causing a loss of PPIX’s intrinsic fluorescence^41, 42^ and PPB readily quenched albumin fluorescence with a dose-response comparable to PPIX indicating the attached biotin moiety (**Fig. 1b**) does not obstruct the binding (**Fig. 1e**). Importantly, biotin alone did not quench albumin fluorescence, revealing the specificity of the PPB-albumin interaction. Additionally, when PPB bound to albumin was passed through a Sephadex G-25 size-exclusion column, PPB alone was retained, but not the PPB:albumin complex, indicating that the majority of PPB was bound to the albumin protein. (**Supplemental Data Fig. 3b-d**). The PPB probe was further developed for application to cells, using cell lysates prepared from the glioblastoma cell line, U251, which were incubated with PPB. In detail, lysates from U251 cells were incubated with either streptavidin beads, PPIX, PPB, or PPB and biotin or PEG biotin **(Fig 2)**. The precipitated pellets were washed 6 times successively in 10 volumes each oof PBS containing 0.1% TX100. Photographs were made of the pellets as a record to monitor the color due to associated porphyrins occurring with cumulative washes (**Fig. 2b**). While lysates incubated with excess PPIX alone prior to interaction with the beads yielded a dark purple bead pellet, commensurate with PPIX amount, this coloration strongly dissipated after the six washes, indicating only weak PPIX association with the beads. Beads exposed to lysates containing both PPB and PPIX showed a strong purple hue which, although becoming lighter during the washing process, retained a color both more intense and different in shade to that seen in beads pulling down with PPB alone. This indicates a strong association of PPIX with the PPB bound beads as suggested in the model (**Fig 2a**, labeled “PPIX competition”). Beads incubated with biotin and PPB showed essentially no coloration, likely due to the failure of PPB to bind. The beads from lysates incubated with the colorless PEG-biotin showed no coloration. Silver staining of the proteins extracted from the pelleted beads with Laemmli buffer (**Fig 2c**) showed that biotin competition was superior to PPIX competition in reducing the pulldown of proteins with PPB, while incubation with PEG-biotin, which is identical to PPB except lacking the PPIX moiety, as expected, yielded only background levels. We replicated this experiment and extended it by using mass spectrophotometry to quantify the eluted proteins and obtained quantitative results similar to what was observed (**Fig 2b,c)**. The results showed that the PPB with PPIX competition was quantitatively similar to PPB alone. Importantly, biotin competition of PPB strongly reduced the number of associated proteins by over 95% (**Fig 2d**). For this reason, biotin competition and PEG-biotin pulldowns were chosen as controls throughout the subsequent experiments.

### Identification of a distinct pattern of PPIX binding in NIH3T3

To identify PPB interacting proteins, PPB-bound protein was precipitated from solubilized whole-cell extracts from four distinct cell lines (**Fig 1A**). To broadly identify PPB binding proteins, the following cell lines of different species and lineage were investigated: MEL, a mouse erythroblast cell line; NIH3T3, a mouse fibroblast cell line; HepG2, a human liver cell line; and U251, a human glioblastoma cell using spectral counts to qualitatively identify the proteins (**Fig. 1A**).

We identified similar proteins among species and lineage by performing quantitative mass spectrometry (MS) analyses of PPB bound proteins from the four cell lines using tandem mass tag (TMT) analysis^43^. Among the PPB-bound proteins, we noted a recently reported novel PPIX binding protein, elongation factor alpha, EEF1A1^22^ (**Supplemental data Full data Table**). We used Significance Analysis of Interaction (SAINT) software for scoring proteins interacting with PPB (**Supplemental Data Saint analysis**). For each cell line, three independent cell lysates were prepared, incubated with PPB, and controls: PEG-biotin alone, and PPB with excess PEG-biotin. Sample loading bias was corrected by using species-specific peptides and PPB interacting proteins determined by SAINT analysis (see **Methods**), allowing for accurate quantification. By comparing with controls, 543 discrete candidates were identified across species as PPB interaction proteins (FDR < 0.01 and z-score > 3**; Supplemental data TMT**). These PPB interacting protein candidates localized to multiple subcellular compartments, as determined by GO analysis, with the largest portion (35%) being cytosolic (**Fig. 1B).** Some of these proteins are likely derived from protein complexes, such as SDHA and SDHB (**Supplemental Data Fig. 4**). Principal component analysis and unsupervised clustering of these proteins revealed that Mel, HepG2, and U251 cells segregated together, but not NIH3T3 cells (**Fig. 1C,D**). Multiple pathway analysis programs and gene ontology databases (KEGG, GO, and Hallmark) were interrogated to identify pathways containing PPB-binding proteins **(Supplemental Data Pathways**). PPB-binding, among human and mouse, proteins were identified in pathways relevant to mitochondria, oxidative phosphorylation, and reactive oxygen species.

**Fig 4.**
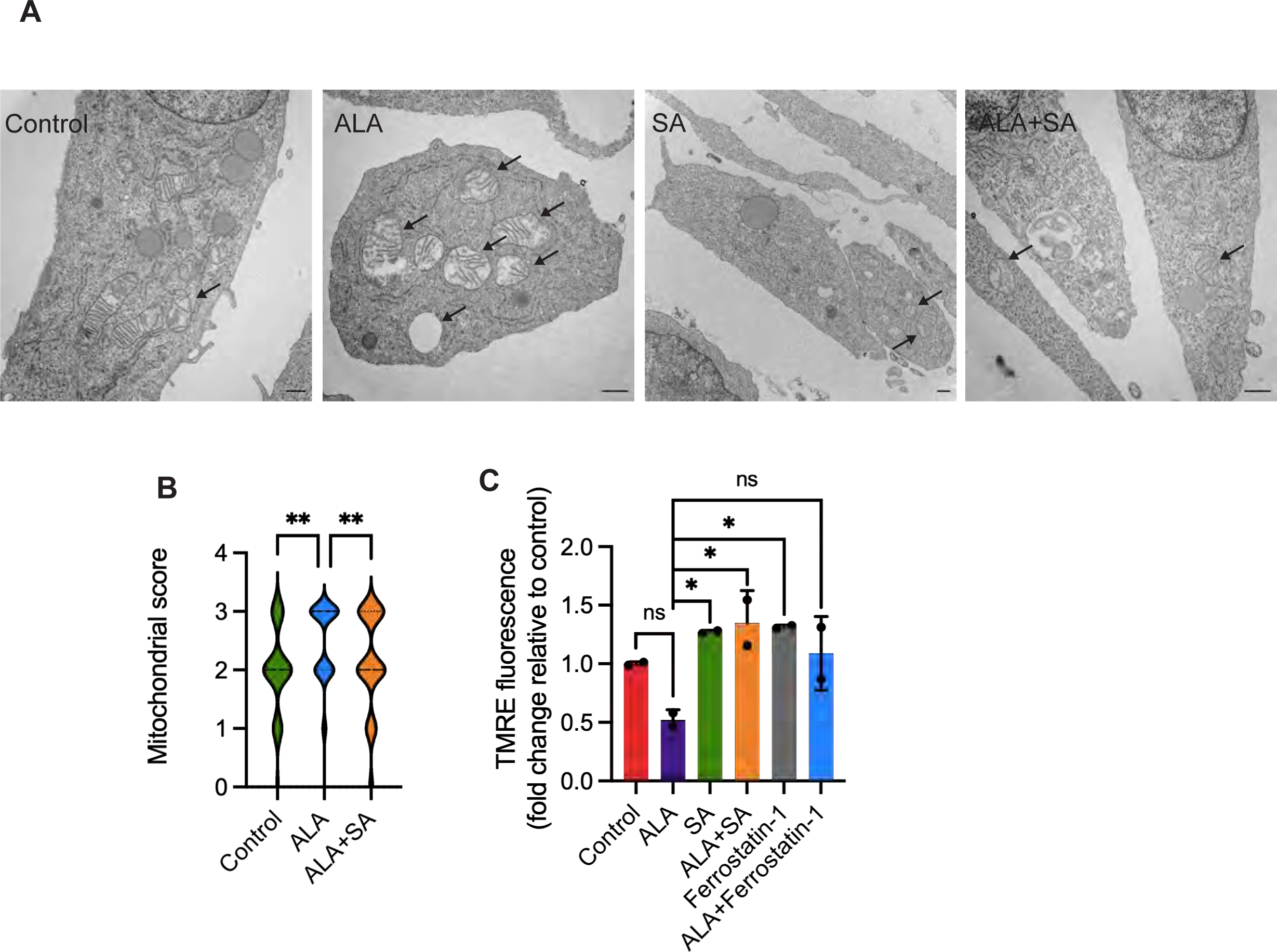
Increased heme/porphyrin synthesis produces altered mitochondria in NIH3T3 cells. **A,** Electron micrographs of representative mitochondria showing morphological abnormalities incurred by ALA treatment (24 hours). Arrows indicate mitochondria showing a score of 2-3 in representative slides. **B,**Average scores of mitochondrial morphology change after ALA treatment(illustrated by violin plot where thickness represents distribution); mitochondria viewed by electron microscopy at 6 hours were scored with 0 indicating unaltered morphology and 3 indicating severe,dose-dependent impairment.(number of mitochondria evaluated were: n=71for Control, n=88 for ALA and n=108 for ALA+SA) **C,** Mitochondrial membrane potential changes with ALA treatment (mean ± SEM, n=2 samples representative of 3 experiments). Significance is indicated by the number of attached asterisks. \**P* < 0.05, \*\**P* < 0.01, ns indicates no significance. Significance in Figure4Awasevaluated using one-way ANOVA, while unpaired Student’s *t*-test, two-sided, was used to determine significance in Figure 4B.

An unweighted gene-clustering analysis^44^ strategy revealed patterns among the PPB-binding proteins that produced eight distinct clusters^45^ **(Supplemental Data Fig. 5)**. For many of these clusters, common proteins were found among the various cell lines. One cluster (cluster 1) appeared to segregate NIH3T3 cells from the other cell lines **(Fig. 4a and Supplemental Data Fig. 5)**.

**Fig 5.**
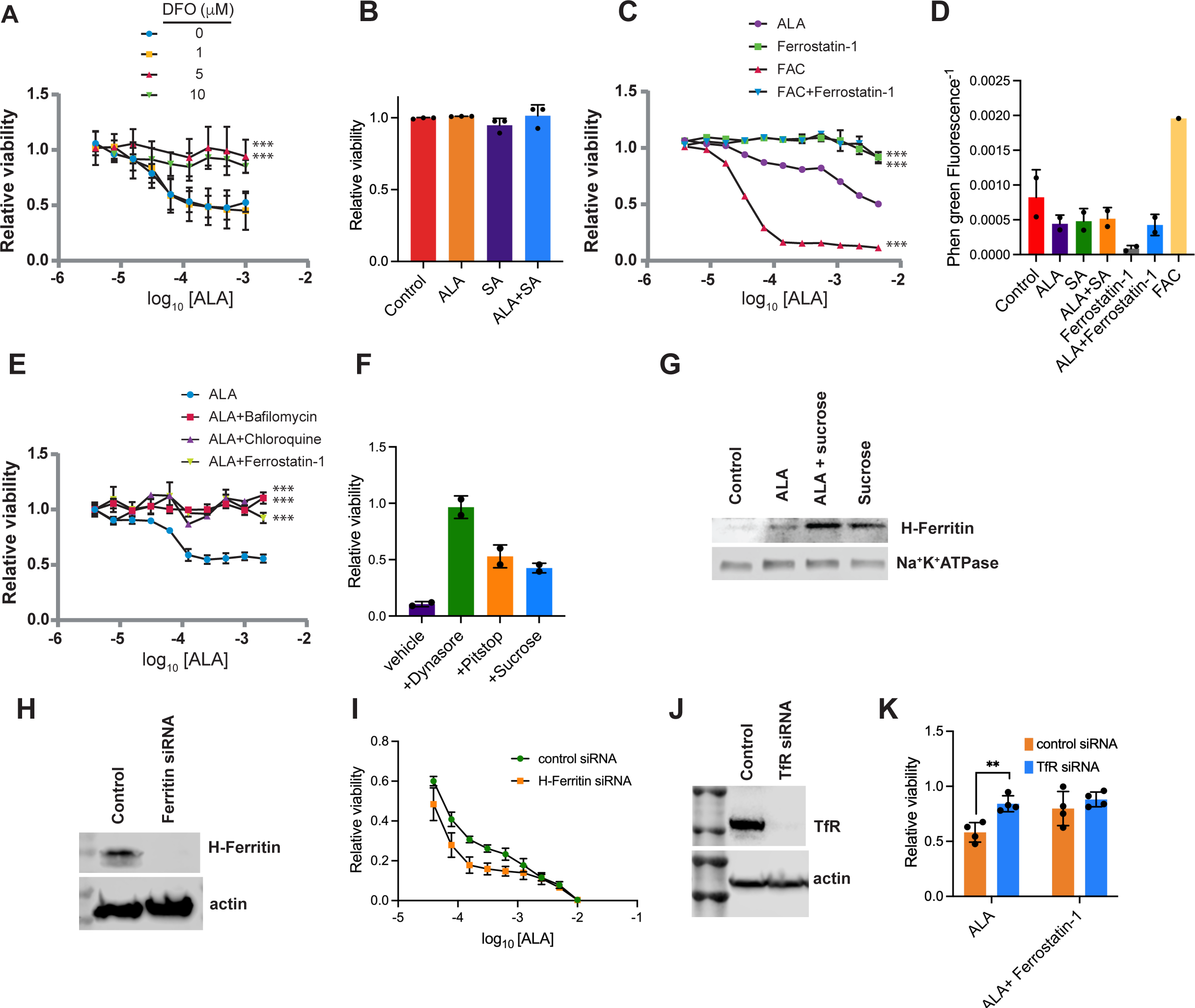
Iron uptake modulates heme/porphyrin synthesis cytotoxicity in NIH3T3 cells. **A,** Desiferrioxamine iron chelation dose-dependently reduces ALA toxicity (mean ± SEM, n=3 independent experiments). **B,** FAC alone produces minimal cytotoxicity (mean ± SEM, n=3 independent experiments). **C,** FAC potentiation of ALA toxicity (mean ± SEM) and abrogation by ferrostatin1(n=2 samples, representative of 3 independent experiments). **D,** ALA treatment does not result in measurable free intracellular iron change (mean of inverse fluorescence ± SEM) as determined by Phen Green fluorescence. (n=2 samples, representative of 3 independent experiments) **E,** ALA cytotoxicity (mean ± SEM) is reversed by both chloroquine (50 µM) and bafilomycin (20 nM) (n=2 samples, representative of 3 independent experiments). **F,** Rescue of ALA-mediated toxicity (mean ± SEM) by clathrin-dependent endocytosis inhibitor Dynasore (15 µM), Pitstop (10 µM) and sucrose (100 µM). (n=2 independent experiments,). **G,** ALA treatment induces movement of ferritin to the membrane related to heme synthesis, leading to initiation and enhancement of ferroptosis. **H,** immunoblot analysis of H-Ferritin expression treated with either control siRNA or specific H-Ferritin siRNA. **I**, Gene knockdown of H-Ferritin sensitizes cells to ALA cytotoxicity(mean ± SEM, n=2 independent experiments). **J**, immunoblot analysis of TfR expression after TfR siRNA knockdown or control siRNA. **K**, Knockdown of TfR protects against ALA cytotoxicity (mean ± SEM, n=4 independent experiments). Significance is indicated by the number of attached asterisks. \**P* < 0.05, \*\**P* < 0.01, \*\*\**P* < 0.001, and ns indicates no significance. Significance in Figure 5B, D, and K were evaluated using one-way ANOVA, while paired Student’s *t*-test, two-sided, was used to determine significance between curves in Figures 5A, C, E, and F.

Several of the most common PPB-interacting proteins in the NIH3T3-specific cluster were related to heme and iron metabolism, including ferrochelatase (FECH), ABCB7, SLC25A4, and SLC25A5, (**Fig. 2B**). By applying String (V11) and Cytoscape network analysis (www.cytoscape.org;V3.7.2), protein inter-relationships connecting heme and iron metabolism were revealed (**Fig 2C, Supplemental Data Table Enrichment genes**). Hypothesizing that relationships between these PPB-binding proteins might lead to the discovery of new relationshipsbetween heme/porphyrin biosynthesis and iron metabolism, we extended our analysis to all eight clusters and proteins bound by PPB in all four cell types. This extension allowed for identification of an array of proteins linked to the ferroptosis pathway (**Fig. 2D**). To our knowledge, a relationship between ferroptosis and an elevation in heme/porphyrin biosynthesis was not known.

### Increasing heme/porphyrin synthesis and ferroptosis

Based on the PPB-binding proteins and network interaction analysis, we hypothesized that increased heme/porphyrin biosynthesis might promote ferroptosis related to PPIX formation. We hypothesized that increased heme/porphyrin synthesis, represented by PPIX elevation, might promote formation of reactive oxygen species (ROS). Heme/porphyrin biosynthesis was activated in NIH3T3 cells by adding the heme precursor aminolevulinic acid (ALA), because it is formed after the rate-limiting step in heme/porphyrin synthesis (i.e., ALAD) and ALA generates both dose- and time-dependent increases in heme and PPIX (**Fig. 3A,B**). We also demonstrated that PPIX accumulates outside the cell in the media in a dose- and time dependent fashion during ALA incubation (**Fig 3C**). To characterize general changes in cellular ROS in ALA-treated cells, dichlorodihydrofluorescein diacetate (H2DCFDA), was used to detect a broad range of ROS (e.g., peroxides and hydroxyl radicals). The superoxide-specific mitochondrial probe, MitoSOX Red was used to determine mitochondrial ROS. ALA treatment increased both mitochondrial superoxide ROS (**Fig. 3D**) and intracellular peroxides **(Fig. 3E)** and required the increase in heme/porphyrin synthesis because succinyl acetone (SA), which blocks porphyrin synthesis through ALAD inhibition, blocked the increase in ROS (**Fig. 3E**). We then investigated whether ALA-generated ROS were associated with loss of viability, and if so, could we rescue the cells. NIH3T3 cells were treated with ALA alone or in combination with the antioxidants ascorbate (Asc) and N-acetyl-cysteine (NAC), which are important for regeneration of reduced glutathione, as well as ferrostatin-1, an antioxidant inhibitor of lipid peroxidation by scavenging alkoyl peroxides^46^ (**Fig 3F**). Succinylacetone (SA), was employed as a negative control. Elevated porphyrin synthesis with ALA treatment strongly reduced viability, an effect blocked or strongly reduced by either inhibition of porphyrin biosynthesis, blocking lipid peroxidation (ferrostatin-1), or supplementation with molecules that promote formation of the antioxidant glutathione (**Fig 3F**).

To elucidate the mechanism of toxicity in ALA-induced porphyrin biosynthesis cytotoxicity, cells were co-treated with one of the following four compounds: ferrostatin-1, which blocks ferroptosis by preventing lipid peroxide formation^46^; necrosulfonamide, which inhibits pyroptosis by binding gasdermin D^47^; GSK-872, a RIP3 kinase inhibitor that blocks necroptosis^48^; or ZVAD-FMK, a pan-caspase inhibitor of apoptosis^49^. ALA alone reduced cell viability in a dose-dependent manner (**Fig. 3G**). Only Ferrostatin-1 suppressed ALA-induced cell death, indicating that increased heme/porphyrin synthesis promotes ferroptosis-related cell death; this cell death is unrelated to apoptosis, necroptosis, or pyroptosis. No alteration in cell cycle accompanied the increased heme/porphyrin synthesis **(Supplemental Data Fig. 6)**.

**Fig 6.**
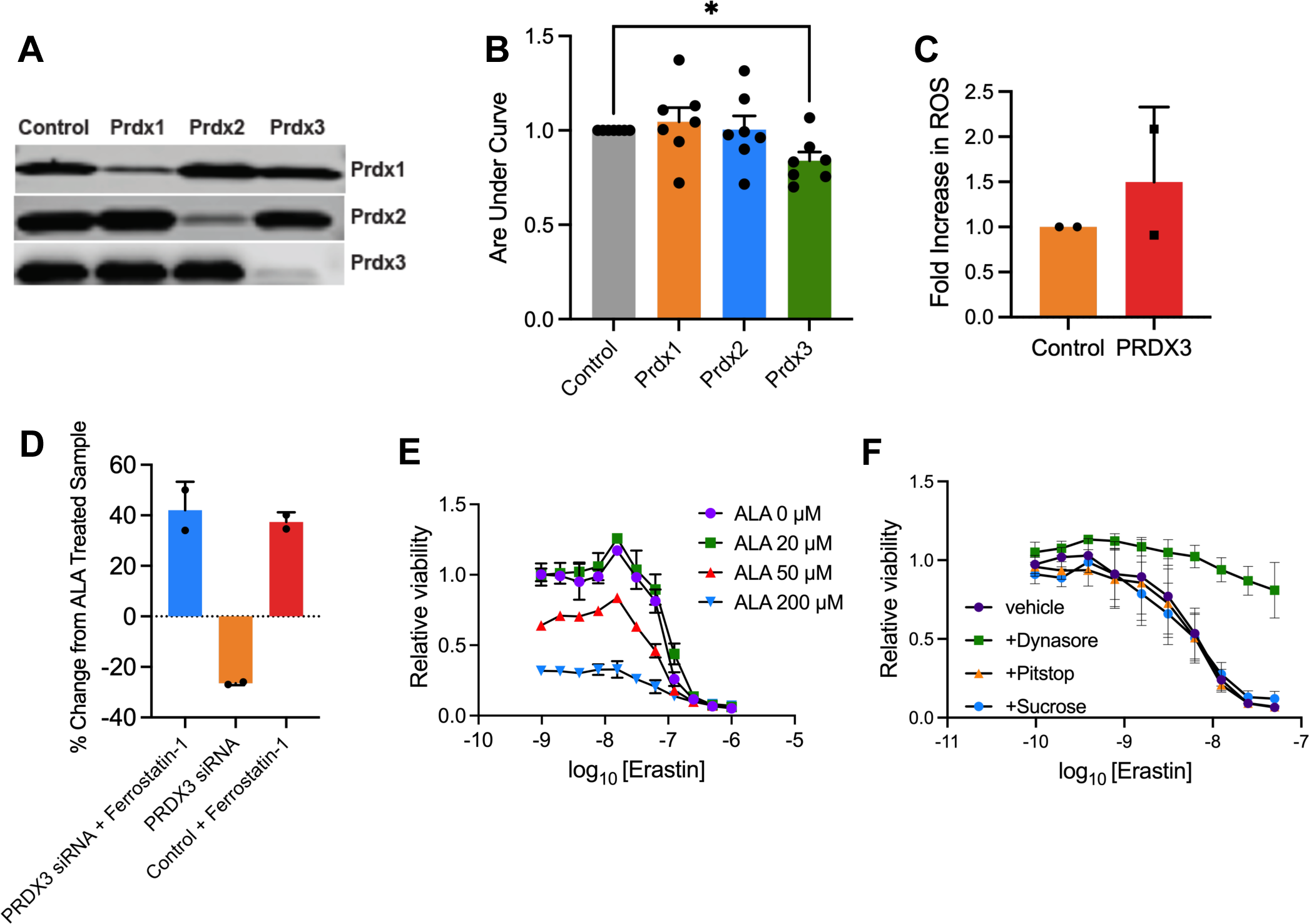
PRDX3 knockdown increases ALA mediated Ferroptosis. **A**, PRDX1, 2,and 3 are efficiently knocked down by siRNA treatment. **B,** Knockdown of PRDX3, but not that of PRDX1 or PRDX2, enhances the cell death (mean ± SEM) caused by PPIX induction (n=6 independent experiments). **C,** Knockdown of PRDX3 increased ROS(mean ± SEM)in both mitochondria (MitoSOX Red) and cytosol (DCF) (n=2 independent experiments). **D**, Treatment with ALA (156μM) induced cytotoxicity(mean ± SEM) that was reversed by Ferrostatin-1, but potentiated by siRNA PRDX3 knockdown, which was reversed by the addition of Ferrostatin-1 (n=2 independent experiments). **E**, Co-treatment of cells with ALA and various concentrations of Erastin increased cytotoxicity (mean ± SEM, n=2 independent experiments,)). P-value is directly indicated in Figure 5B for Control vs. PRDX3.

Death from ferroptosis is tightly coupled to lipid peroxide formation in cellular membranes^50^. To confirm whether lipid oxidation occurs within the context of ALA-promoted porphyrin synthesis, we used the BOPIDY-C11 probe. This fluorescent probe intercalates into membranes, permitting peroxidation of the surrounding lipid moieties to alter its fluorescence. Induction of porphyrin biosynthesis promoted increased BOPIDY-C11 fluorescence (**Fig. 3H**), that was abrogated by SA treatment showing that increased porphyrin biosynthesis, but not ALA alone which is incapable of promoting lipid peroxidation. The antioxidant enzyme GPX4, a gluthione peroxidase, protects against peroxide damage to the cell by oxidizing glutathione co-factor to defend against ferroptosis^51^. Oxidized glutathione levels strongly increased after ALA treatment (**Fig. 3I**), thus revealing that sustained heme/porphyrin biosynthesis is an oxidative challenge that impacts the regeneration of reduced glutathione. We next determined whether GPX4 activity was increased in cells that displayed activation of heme/porphyrin synthesis by ALA. We show that GPX4 activity, as measured by an assay coupled with glutathione reductase, was increased by over two fold **(Supplemental Fig. 9)**, consistent with the increased C11-BODIPY peroxidation.

Using transmission electron microscopy, we investigated whether increased levels of ALA-induced l ROS caused mitochondrial changes. We discovered that NIH3T3 cells treated with ALA, had strongly increased the proportion of cells with mitochondrial morphology and that these morphological changes were blocked by inhibition of porphyrin biosynthesis with SA (**Fig. 4A and B**). The mitochondria showed increased deformity and cristae loss (**Fig. 4A and B**). Given the evidence of mitochondrial injury and the knowledge that PPB binds ATP translocase, we investigated whether mitochondrial membrane potential was altered by the elevation in PPIX associated with the increased heme/porphyrin biosynthesis by using the polarity-sensitive mitochondrial dye, TMRE. The mitochondria showed strong depolarization with ALA treatment, consistent with both ATP translocase inhibition and ROS generation. The mitochondrial depolarization was rescued by both porphyrin synthesis inhibition with SA and ferroptosis inhibition (**Fig. 4C**)

### Iron modulation of heme/porphyrin biosynthesis ferroptosis

Ferroptosis is an iron-dependent, non-apoptotic death^52^. To assess the role of intracellular iron in the PPIX-induced viability loss, we modulated intracellular iron concentrations in the presence of elevated porphyrin/heme synthesis. To reduce the free intracellular iron levels, we treated cells with the iron chelator desferrioxamine (DFO), which dose-dependently suppressed ALA cytotoxicity **(Fig. 5A**). DFO can induce toxicity but was only mildly toxic in this study’s conditions **(Supplemental Data Fig. 8)**. Conversely, intracellular iron concentrations were elevated by treatment with the cell-penetrant form of iron, ferric ammonium citrate (FAC). FAC treatment alone was non-toxic **(Fig. 5**) but strongly enhanced the cytotoxicity of ALA in a manner inhibited by the ferroptosis inhibitor ferrostatin-1 (**Fig. 7c**). Thus, porphyrin/heme synthesis-induced cell death, during ALA treatment, is potentiated by iron augmentation and attenuated by its sequestration. Consequently, we sought to determine whether PPIX initiates ferroptotosis by directly increasing intracellular free iron. The fluorescent iron-sensor Phen-Green was used to determine whether increased heme/porphyrin synthesis by ALA is associated with measurable, free intracellular iron change. Interestingly, there was no evidence fora detectable change in free-iron by Phen-Green **(Fig. 5D).**

**Fig 7.**
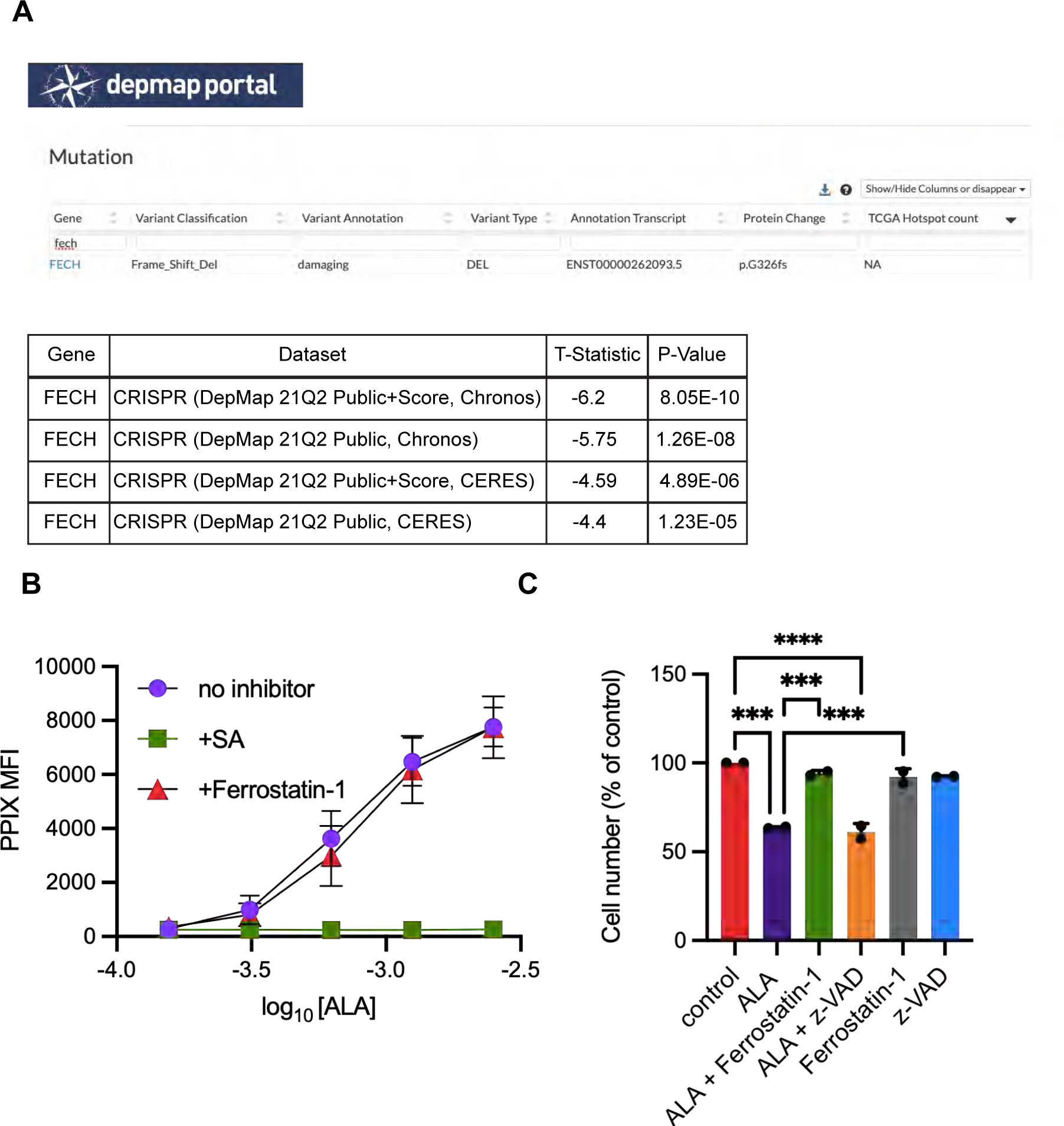
Sensitivity of Jurkat cells with defective Fech to elevated porphyrin synthesis. **A**, Jurkat T-lymphoblastoid cells harbor deletion in the *Fech* gene producing a frameshift and deletion. CRISPR mediated deletion of Fech reveals among multiple datasets with significant dependency (source Depmap (depmap.org/portal). **B**, Various ALA concentrations were used to treat Jurkat cells followed by analysis of PPIX concentration (mean ± SEM). Cells were treated with either ALA alone, ALA+Ferrostatin-1 to assess whether it affected PPIX formation of ALA, plus succinyl acetone (SA) (n=2 samples, representative of 3 independent experiments). C, Viability following treatment with either ALA, ALA and ferrostatin-1, ferrostatin-1 alone or ALA and ZVAD-FMK. Significance is indicated by the number of attached asterisks. \**P* < 0.05, \*\**P* < 0.01, \*\*\**P* < 0.001 ****P<.0001, and ns indicates no significance. Significance in Figure 6C was evaluated using one-way ANOVA.

To further investigate the role of iron in ALA-promoted cell death, we investigated modes of iron uptake. Iron can be chaperoned into the cell by the transferrin receptor (**TfR1**) or by H-Ferritin^53, 54^ and subsequently released after fusing to the lysosome^34^. It is sequestered in a complex formed from H-ferritin and L-ferritin^55^. We reasoned that inhibitors of lysosome function, chloroquine and bafilomycin, might restrict lysosomal iron uptake. Both agents, at concentrations that were not toxic were as effective as ferrostatin-1 in attenuating porphyrin synthesis–induced cell death (**Fig. 5E**). Additionally, we assessed the role of clathrin-dependent endocytosis using either the dynamin inhibitor Dynasore (DS), sucrose or Pitstop 2. We used these agents because of Dynasore’s reported off-target effects^56–58^. Dynasore strongly protected against ALA-promoted ferroptosis. Both sucrose and Pitstop provided protection too (**Fig. 5F**). To determine whether porphyrin/heme synthesis status affected membrane-localized H-ferritin, we employed a membrane surface biotinylation–labeling approach^59^. We assessed whether increased heme/porphyrin synthesis induced by ALA resulted in a greater amount of H-ferritin at the membrane. Interestingly, the combination of ALA treatment with blockade of endocytosis by either Dynasore or sucrose treatment increased the amount of H-ferritin at the membrane (**Fig. 5G)**, suggesting that ALA treatment promoted membrane localization of H-ferritin. Using H-Ferritin siRNA knockdown (**Fig. 5H**), the role of H-Ferritin in either sequestration or of shuttling iron into the cell was assessed under conditions where ALA-activated heme/porphyrin synthesis. ALA-cytotoxicity was enhanced by H-Ferritin knockdown over a broad range of ALA concentrations. This suggests that sequestration of intracellular iron, rather than import, is the primary function of H-Ferritin in heme/porphyrin synthesis induced ferroptosis (**Fig. 5I**).

Transferrin receptor (TfR) mediated uptake of iron begins with the capture of iron-laden transferrin by the receptor (TfR), initiating internalization and formation of endosomes^60^. These endosomes, now harboring iron, subsequently release iron intracellularly. We interrogated the role of TfR in heme/porphyrin synthesis mediated ferroptosis. Surface membrane expression of TfR was unaltered by ALA-increased porphyrin biosynthesis. However, knockdown of TfR (**Fig. 5J**) reduced the ALA-induced cytotoxicity to a similar degree as did Ferrostatin-1(**Fig. 5K**). In total, we interpret these findings to indicate that both H-Ferritin and the TfR are important in the importation of extracellular iron during ALA induced porphyrin biosynthesis, however, overall, they have divergent functions.

### PRDX3, a PPB interactor, modulates heme/porphyrin ferroptosis

Given that ALA elevated ROS and that exogenous antioxidants protected against ALA-induced toxicity, we queried whether modulation of antioxidant enzymes contributed to the ferroptotic process. We noted that three of the six peroxiredoxins (PRDXs), PRDX1, PRXD2, and PRDX3 bound PPB. These proteins comprise ubiquitous cysteine-dependent anti-oxidant enzymes that protect against peroxides by metabolizing hydrogen peroxide^61^. We tested whether PRDX1-3 protected against ALA-promoted ferroptosis by suppressing PRDX 1-3 using siRNAs and then determining how ALA cytotoxicity was impacted. First, we verified that each siRNA suppressed its respective PRDX family member (**Fig. 6A**). Cytotoxicity of ALA over a range of ALA concentrations was significantly increased when only PRDX3 was suppressed with siRNA (**Fig. 6B**), indicating that this protein helps modulate heme/porphyrin synthesis–mediated ferroptosis. PRDX3 reduction also increased cytosolic ROS, a response secondary to activation of heme/porphyrin synthesis (**Fig. 6C**). To confirm that PRDX3 knockdown impacts ALA-mediated ferroptosis, cells were treated with either control siRNA or the siRNA to PDRX3 followed by either ferrostatin-1, ALA alone, or ALA plus ferrostatin-1. Cells treated with PRDX3 siRNA and ALA had a 27% loss in viability relative to treatment with ALA alone. Ferrostatin-1 increased viability (42%) in cells treated with both PRDX3 siRNA and ALA (**Fig. 6D**), indicating that PRDX3 has an important role in modulating porphyrin synthesis induced ferroptosis.

To further delineate the elements of porphyrin synthesis-induced ferroptosis, we investigated whether cell death initiated with the classic ferroptosis inducer, erastin, could be affected by the state of heme/porphyrin synthesis. Increasing heme/porphyrin synthesis sensitized cells to erastin (**Fig 6E**), and a pharmacological interaction analysis showed a negative alpha value (-0.707, p=1.25e-16) indicating synergism (**Supplemental Data Fig 11**). Suppression of heme synthesis by succinyl acetone (SA) did not protect against erastin toxicity (**Supplemental Data Fig. 12**). SA readily reversed ALA-induced cytotoxicity, but was incapable of affecting cell death initiated by erastin, a cystine/glutamate antiporter inhibitor. Moreover, the lowest concentration of ALA had minimal effect of erastin toxicity, likely due to the small elevation in intracellular PPIX, while ALA concentrations of 50 and 200 μM strongly enhanced cytotoxicity (**Fig. 6E**). We also show that neither Pitstop nor sucrose affected erastin toxicity which further underscores that heme/porphyrin biosynthesis driven death is separable from canonical ferroptosis (**Fig 6F**).

### Heme/porphyrin biosynthesis, ferroptosis and a lymphoblastic leukemia cell line

Because of the relationship between PPIX formation with activation of heme/porphyrin biosynthesis, we interrogated the cancer dependency map for cell lines that might rely on the terminal enzyme in heme biosynthesis, FECH. According to the DepMap (Depmap.org), many, though not all, cancer cell lines are highly reliant on the terminal enzyme in heme biosynthesis, FECH. We hypothesized that cells deficient in Fech, unable to efficiently convert PPIX to heme, might be susceptible to ferroptotic death promoted by elevated porphyrin biosynthesis. Interrogating the DepMap, we identified the T-lymphoblastoid cancer cell line, Jurkat, as a *FECH-*deficient cell line due to a frame shift deletion in *FECH* (**Fig. 7A**). Next, we investigated whether increased porphyrin synthesis by ALA treatment produced elevated intracellular PPIX in Jurkat cells. Using FACS analysis, we found ALA dose-dependently increased the intracellular PPIX (**Fig. 7B**). We then evaluated whether elevated PPIX promoted cell death in Jurkat cells, and whether that death occurred by ferroptosis. Cells were treated with either ALA, ALA and ferrostatin-1, ferrostatin-1 alone, or ALA and ZVAD-FMK, the pan-caspase inhibitor **(Fig. 7C)**. Cell viability was reduced by almost 50% by ALA treatment and viability was not restored by the addition of ZVAD-FMK. In contrast, ferrostatin-1 was capable of rescuing cell death produced by ALA treatment. These findings suggest cancer cells susceptible to PPIX elevation, such as those deficient in FECH, might be killed by increasing PPIX secondary to elevated porphyrin biosynthesis.

## Discussion

Previously, probes such as hemin coupled to agarose beads had been employed to determine the repertoire of heme-binding proteins in particular cells or tissues^62^. However, PPIX, a prominent tetrapyrrole intermediate of the heme biosynthetic pathway, has never been used to probe for protein-protein interactions^63^. Here we describe for the first time, the development, characterization and use of a PPIX probe (PPB) to interrogate the cellular proteins that interact with protoporphyrin IX in a variety of cell types. Lysates from cell lines, NIH3T3, HepG2, MEL and U251, representing four different histotypes (two from each, mouse and human) with diverse capacity for heme synthesis, were analyzed for PPB binding with the expectation that the PPIX interactome might differ between them, providing additional insight. For example, HepG2 hepatoblastoma cells are known to express heme-requiring P450 enzymes^64^, MEL cells are able to generate large amounts of hemoglobin upon differentiation, and ALA treated U251 glioblastoma cells are known to accumulate high levels of PPIX^65, 66^. We discovered an array of proteins that, surprisingly, were common among these cells but distributed throughout multiple subcellular compartments (plasma membrane, cytosol, mitochondria, endoplasmic reticulum, Golgi, and nuclear envelope). This finding was consistent with PPIX’s broadly distributed intracellular immunofluorescence and the subcellular localization of PPIX binding proteins^25, 27–33, 51, 67, 68^. Through curation and pathway analysis of these proteins, we discovered an unanticipated relationship between ferroptosis and heme biosynthesis. Additionally, we were able to identify a relationship between this novel ferroptosis pathway and a PPB-interacting protein, PRDX3. PRDX3 is a mitochondrial ROS scavenger^69^ that mitigates the cell death promoted by activation of the heme biosynthesis pathway.

Previous studies have investigated cellular heme-binding proteins through their binding to hemin-bound agarose beads^62, 70^. To date, no such comparable effort to broadly identify proteins interacting with the penultimate, non-iron, precursor of heme, PPIX, has been reported. Our strategy involved a two-stage approach, allowing cellular proteins to first interact with PPB, followed by PPB bound proteins captured with avidin affinity beads. A likely advantage for this strategy over conventional agarose-coupled probes (e.g. hemin-agarose) was potentially facilitating interaction by reducing steric hindrance. Isolating the probe-protein complexes was easily accomplished using commercially available streptavidin beads, enabling convenient identification. Furthermore, by including an excess biotin control, background binding is eliminated to improve detection. Attempts to use excess PPIX as a competitor for non-specific binding failed, possibly due to excessive background proteins associated with the high concentration of PPIX needed to compete.

Our procedure for identifying cellular PPB-binding proteins appears robust, as we successfully identified a previously reported PPIX-binding proteins, such as EEF2A1^29^ and ferrochelatase^13^, among the PPB-bound proteins. In addition, some of the PPB-bound proteins bind heme, suggesting the common feature of recognition is the tetrapyrrole (see **Supplemental Data Fig 10**). It is extremely unlikely, however, that any of the results obtained were due to conversion of the PPIX moiety to a heme moiety, as ferrous iron, required by enzymatic incorporation via ferrochelatase, requires near anaerobic conditions^71^, and this anaerobic environment was not attainable during our room temperature incubations. It is possible, for common heme- and PPIX-binding proteins, that PPB may displace heme. Interestingly, ABCG2, a plasma membrane transporter known to bind and export PPIX from the cell, was not detected. The inability to detect ABCG2 could be due to either the extraction conditions or, perhaps the level of ABCG2 in these cells was too low to be detected.

We noted by PCA analysis, an unbiased technique for visualizing differences in data sets, that NIH3T3 cells were distinguishable from the other three cell lines. Cluster analysis also revealed a well-defined set of proteins that was more abundant in the NIH3T3 cells. Many of these PPB-associated proteins were related to iron and heme synthesis, including MMS19^72^ ferrochelatase^73^, ABCB7^74^, and both SLC25A4 and SLC25A5, which form the mitochondrial ADP/ATP translocase^75^ and have been shown to import PPIX and heme^76^. We investigated the NIH3T3 cells for the impact of elevated heme/porphyrin synthesisreflected by increased PPIX when incubated with ALA.

ALA induction of heme/porphyrin synthesis in NIH3T3 cells revealed a susceptibility to cell death that was not evident in liver derived HepG2 cells. Interestingly, susceptibility also differed greatly with respect to the fetal calf serum used to culture the NIH3T3 cells, possibly due to variation of some unknown serum component. Chemical inhibitors of multiple forms of cell death, such as apoptosis, pyroptosis, or necroptosis, were incapable of preventing heme/porphyrin synthesis–induced cell death. Cell death was, however, readily blocked by succinylacetone, which blocks heme/porphyrin synthesis at ALA-dehydratase, whose substrate is ALA. Significantly, porphyrin-mediated cell death could also be readily reversed by ferrostatin-1, a ferroptosis inhibitor that blocks lipid peroxidation^10, 77^ as well as prototypical scavengers of ROS such as N-acetyl cysteine.

Importantly, loss of viability was preceded by increased ROS levels, glutathione depletion, and lipid peroxidation, features that closely align with the canonical features of ferroptosis^50, 78, 79^. We found that PPIX was dose- and time-dependently exported from NIH3T3 cells and hypothesized that exogenous PPIX might also elicit cell death. Exogenous treatment of NIH3T3 with PPIX, but not heme, generated dose-dependent decreases in cell viability suggesting exported PPIX could be lethal to adjacent cells. Interestingly, heme/porphyrin synthesis–induced cell death was preceded by elevated ROS, impairment of mitochondrial integrity and mitochondrial depolarization, which contrasts with some ferroptotic systems where hyperpolarization of mitochondria is observed^80, 81^. Together this data strongly supports a model in which ALA-induced heme/porphyrin synthesis produces a form of ferroptosis in NIH3T3 that is distinct.

Iron uptake and iron trafficking contribute to intracellular iron homeostasis, an important aspect of ferroptosis^81–83^. For cell death induced by elevated porphyrin synthesis, intracellular concentrations of iron and its transport appear to be important factors. Exogenous iron enhanced ALA induced cytotoxicity, while intracellular iron chelation with desferrioxamine (DFO) reduced cell death, supporting the role of intracellular iron in heme/porphyrin synthesis–induced cytotoxicity. The lysosomal inhibitors bafilomycin and chloroquine also blocked heme/porphyrin synthesis-induced cell death, implying that iron trafficking through the lysosome is a key component of heme/porphyrin synthesis– induced cell death. Treatment with inhibitors of endocytosis such as the dynamin inhibitor, Dynasore, sucrose and Pitstop thwarted heme/porphyrin synthesis–induced cell death. Dynasore has been demonstrated to block ferroptosis through diverse pathways, not only by inhibiting transferrin receptor endocytosis, but also by mitigating ROS by affecting mitochondrial respiration^58^. Consequently, Dynasore’s effects must be interpreted with caution. However, inhibition of endocytosis with hypertonic sucrose also blocks death, supporting the view that iron transit into the cell may occur through an endocytic process. Iron is bound and trafficked to the lysozyme by plasma membrane receptors such as transferrin receptor (TfR1) and H-ferritin^82, 83^. Elevated heme/porphyrin synthesis appears to increase H-ferritin levels at the plasma membrane, with greater accumulation when endocytosis is blocked. The role of H-ferritin was not initially clear, however knockdown of H-ferritin enhanced cell death indicating a protective role. Intracellular iron sequestration with DFO blocked cell death also supports the role of H-ferritin and L-ferritin likewise binding intracellular iron to prevent its release. We also show that siRNA-mediated knockdown of TfR1 blocked heme/porphyrin synthesis-induced ferroptosis consistent with its previous reported role in iron uptake to promote ferroptosis. Collectively, these studies show that both H-Ferritin and TfR1 have distinct, but separable roles in modulating the ferroptosis induced by increased porphyrin synthesis.

Generation of ROS is central to the mechanism of ferroptosis and has been held in check by antioxidant enzymes like glutathione peroxidase GPX4^84^. Another class of proteins that also scavenge ROS are the ubiquitous peroxiredoxins (PRDX)^85–87^, three of which we found to complex with PPB. We hypothesized that suppression of these enzymes, either alone or in combination, might enhance heme/porphyrin synthesis–induced cell death. However, only PRDX3 silencing enhanced toxicity. As PRDX3 localizes to the mitochondria^87^, this result was surprising, and suggested that PRDX3 expression is needed to suppress heme/porphyrin synthesis-initiated cell death. Interestingly, lipid peroxidation is suppressed by the cytosolic PRDX1 in corneal epithelial cells^88^. Coupled with our findings, this indicates that the PRDXs might function to modulate ROS in a subcellular compartment–dependent fashion.

Our study employed an approach using a novel PPIX-biotin (PPB) probe to capture PPIX-interacting proteins in four cell lines, diverse with respect to histotype, species, and heme/porphyrin synthesis capacity. We then analyzed that data to make predictions about PPIX function, resulting in several key findings. First, through gene set enrichment and pathway analysis, we discovered that activation of the heme/porphyrin biosynthetic pathway promotes ROS formation and death consistent with ferroptosis in NIH3T3 cells. We subsequently accurately predicted that a ferrochelatase deficient cancer cell line, Jurkat, would be similarly susceptible. This implies that other cancer cells, such as those harboring impaired ferrochelatase function might also be vulnerable to this form of cell death. Secondly, we were able to further use our data to predict that an antioxidant enzyme from our screen, PRDX3, might be pertinent to this pathway by showing that porphyrin-induced ferroptotic cell death is moderated by the PPB-binding protein, PRDX3. However, how it is integrated with TfR, H-Ferritin into modulating heme/porphyrin cell death is unknown at this point.

Our discoveries suggest that ferroptosis might be a mechanism contributing to cell death under PPIX elevation due to an alteration of the heme/porphyrin pathway activity related to genetic disease, cancer, or toxin exposure^89, 90^,. These findings suggest that using biotin appended molecules might be applied to identify relationships between such molecules and the biological pathways the impact. Further analysis of our PPB data might also lead to additional insights. For example, intriguingly, several of the highly significant PPB-interacting proteins that were identified in have roles in metabolism-related pathways such as oxidative phosphorylation, ROS, and mTORC1 (see **Supplemental Data Table 5**), suggesting that these PPB proteins might modulate their activity and indicating a potential area for future studies.

## Methods

### Compounds

The following reagents were used: d-aminolevulinic acid, bafilomycin, chloroquine, ferrostatin-1, (Sigma), Laemmli buffer (GeneTex), Bodipy 581/591 C11, Phen Green SK, (Invitrogen), TMRE (Abcam), GSK872, necrosulfonamide, (Cayman), Liproxystatin (Tocris), and Dynasore (Abcam).

### Cell Lines

NIH3T3, K562, and U251 cells were cultured in Dulbecco’s modified Eagle’s medium (Gibco) without nucleosides and supplemented with 10% fetal bovine serum (Hyclone for NIH3T3 cells, Gibco for others), 100 units/mL penicillin/streptomycin and 2 mM L-glutamine. MEL cells were cultured in RPIM 1640 medium (Gibco) without nucleosides and supplemented with 10% fetal bovine serum, 100 units/mL penicillin and 2 mM L-glutamine. HepG2 cells were cultured in Minimum Essential Medium α (Gibco) with 10% fetal bovine serum and 100 units/mL penicillin.

The following antibodies were used where indicated in the text:

| 562 | <u>Name</u> | <u>Source</u> | <u>Catalog#</u> |
| --- | --- | --- | --- |
| 563 | Ferritin | Abcam | AB 75973 |
| 564 | NaK ATPase | Novus | NB 300-146SS |
| 565 | Prdx1 | Invitrogen | LF-MA0073 |
| 566 | Prdx2 | Invitrogen | LF-MA0144 |
| 567 | Prdx3 | Abnova | MAB2725 |
| 568 | TfR | Zymed | 13-6800 |
| 569 |  |  |  |

### Albumin fluorescence quenching

Bovine serum albumin (BSA) fluorescence was determined by using 4 µM albumin in phosphate-buffered saline (PBS). Each compound was prepared in DMSO and then added to 100 µL of 4 mM albumin in PBS in triplicate lanes of a 96-well plate and allowed to bind for 15 minutes. Equivalent volumes of DMSO alone were added to negative control wells. Fluorescence was then assessed at wavelengths of 290-nm excitation and 337-nm emission and normalized to untreated albumin.

### PPIX-biotin (PPB) Probe Synthesis

The Fmoc protecting group was removed from Fmog-PEG Novatag resin (170 mg) using 20% piperidine in dimethylformamide (DMF). The protoporphyrin IX (PPIX) moiety was prepared by combining PPIX (25 mg) with 16.3 mg of 1-Hydroxy-7-azabenzotriazole N,N’-diisopropylcarbodiimide dissolved in 3 mL of N-methyl-2-pyrrolidone, which activated the carboxylic group and formed a symmetrical anhydride. The PPIX moiety mixture was added to the deprotected resin and left for 3 days at room temperature.

### PPB precipitation assays

Assays were performed using 500 µg of lysate per 50 µL of avidin beads. Lysates were first precleared by incubation with 50 µL of beads at room temperature for 30 minutes. Supernatants were transferred to fresh tubes and 40 nmol of PPB or biotin with only a cross-linker as a control were added to the tubes. The tubes were allowed to equilibrate in the dark for 30 minutes. To the cleared lysate, 50 µL of beads in 50% slurry with PBS, 0.1 % Triton X100, was added and incubated an additional 45 minutes in the dark with occasional mixing. The beads were precipitated by centrifugation at 500 x *g* for 2 min and washed 8 times in 10 volumes of PBS/0.1% Triton X100. Proteins were eluted in 2X Laemmli loading buffer with B-mercaptoethanol at 95°C for 5 min before subsequent analysis.

### Protein digestion and peptide isobaric labeling by Tandem Mass Tag (TMT)

Performed as previously described with slight modifications^40^,. First, 2 µg of protein for each sample was electrophoresed on a short gel and stained by Coomassie blue before the protein concentration was determined using BSA as a standard. After protein estimation, gel bands were cut into smaller pieces for in-gel digestion. The gel bands were first washed with 50% acetonitrile, then reduced by adding 5mM DTT at 37°C for 30 min followed by alkylation with 10 mM iodoacetamide (IAA) for 30 min in the dark at room temperature. Unreacted IAA was quenched with 30 mM DTT for 30 min. The gel bands were then washed, dried in a speed vacuum, and rehydrated with a buffer containing trypsin (Promega). Samples were digested overnight at 37°C and acidified using ?. The peptides were extracted in acetonitrile, dried in a speed vacuum, reconstituted in 50 mM HEPES (pH 8.5), and labeled with 11-plex Tandem Mass Tag (TMT) reagents (Thermo Scientific) according to manufacturer’s recommendations.

### Two-dimensional HPLC and Mass Spectrometry

The TMT-labeled samples were mixed equally, desalted, and fractionated over a 60-min gradient on an offline HPLC (Agilent 1220) by using basic pH reverse-phase liquid chromatography (LC). The fractions were dried and resuspended in 5% formic acid and analyzed by acidic pH reverse-phase LC-MS/MS analysis. The peptide samples were loaded on a nanoscale capillary reverse-phase C18 column (New objective, 75 um ID x ∼25 cm, 1.9 µm C18 resin from Dr. Maisch GmbH) by HP and eluted over a 60-min gradient. The eluted peptides were ionized by electrospray ionization and detected by an in-line Orbitrap Fusion mass spectrometer.

The mass spectrometer is operated in data-dependent mode with a survey scan in Orbitrap (60,000 resolution, 2 X 105 AGC target and 50 ms maximal ion time) and MS/MS high-resolution scans (60,000 resolution, 2 X 105 AGC target, 200 ms maximal ion time, 38 HCD normalized collision energy, 1.5 m/z isolation window with 0.2 m/z offset, and 20 s dynamic exclusion).

### Identification and quantification of proteins with JUMP software suite

Proteins were identified and quantified by using the JUMP proteomics software suite to evaluate the false-discovery rate (FDR). All original target protein sequences were reversed to generate a decoy database that was concatenated to the target database. The target protein database was generated by combining both human and mouse protein sequences downloaded from UniProt, with contamination proteins added. Major parameters included precursor and product ion mass tolerance (± 15 ppm), full trypticity, static mass shift for the TMT tags (+229.16293), carbamidomethyl modification of 57.02146 on cysteine, dynamic mass shift for Met oxidation (+15.99491), maximal missed cleavage (n = 2), and maximal modification sites (n = 3). Putative peptide-spectrum matches (PSMs) were filtered by mass accuracy and then grouped by precursor ion charge state and filtered by JUMP-based matching scores (Jscore and ΔJn) to reduce the FDR below 1% for proteins. If one peptide could be generated from multiple homologous proteins, based on the rule of parsimony, then the peptide was assigned to the canonical protein form in the manually curated Swiss-Protein database.

### Tandem Mass Tag-based quantification analysis

TMT-based quantification analysis was performed in the following steps. First, TMT reporter-ion intensities of each PSM were extracted. Second, the raw intensities were corrected based on isotopic distribution of each labeling reagent (e.g., TMT126 generates 91.8%, 7.9% and 0.3% of 126, 127, 128 m/z ions, respectively). Third, PSMs of very low intensities (e.g. minimum intensity of 1,000 and median intensity of 5,000) were excluded. Fourth, the mean-centered intensities were calculated across samples (e.g. relative intensities between each sample and the mean). Fifth, protein or phosphopeptide relative intensities were summarized by averaging related PSMs. And sixth, the relative intensities were multiplied by the grand-mean of the three most highly abundant PSMs to derive protein or phosphopeptide absolute intensities.

### Adjustment of loading bias by species-specific peptides

To minimize technical variation, loading-bias correction has become a crucial step for quantitative proteomics analysis, based on the assumption of equal input amount across all samples. However, this assumption may not be valid for precipitation experiments. To overcome this, we pooled both human and mouse cell lines into one 11-plex TMT experiment so that species-specific peptides were leveraged to correct loading bias across samples, with the following assumption: mouse-specific peptides in human cell lines should have TMT signals similar to those of blank samples, whereas the opposite should be true for human-specific peptides. Briefly, all quantified peptides were separated into three categories: human-specific peptides, mouse-specific peptides, and peptides shared between the two species. To adjust loading bias in a human cell line, we defined the loading-bias normalization factor as median (TMT signal across PSMs from mouse specific peptides in that cell line) / median (TMT signal across PSMs from mouse specific peptides in the 1st blank sample). A similar strategy was implemented to adjust loading bias for mouse cell lines. The corrected data were used for all downstream analyses (e.g., differential expression, network analysis).

### Statistical analysis for identification of PPIX interaction candidates

Four cell lines (two human and two mouse) with duplicates and three blank negative control samples were analyzed. To avoid bias of protein quantification due to species-specific peptides, only homologous peptides shared between human and mouse were used for protein quantification. For each of the four cell lines, interaction candidates were identified by comparison to the three blank samples via one-tailed *t*-test. The four results (i.e., the P value for individual tests) were combined into one integrative P value by using Fisher’s method, and the resulting combined P value was then converted to FDR (for multiple tests correction) by using the BH method. In addition, the measurement variation was based on the replicated measurements of the samples. The relative expression (i.e. ratio) between replicates for each protein was calculated. The ratios of all proteins from the replicates were fitted with a Gaussian distribution to estimate expected mean and standard deviation (null SD). Proteins showing a combined FDR < 0.01 with log2 fold change larger than 3 null SD were considered to be interaction candidates.

### Co-expression and protein-protein interaction (PPI) network analysis

The analysis was performed by using JUMP software as previously described. Briefly, the identified PPIX interaction candidates were clustered into eight co-expression clusters by using the WGCNA R package^45^. Proteins in each cluster were then superimposed onto a composite PPI database by combining STRING-identified PPI data (STRING (v10) STRING v10: protein-protein interaction networks, integrated over the tree of life). Modules in each protein cluster were defined by calculating a topologically overlapping matrix for the PPI network and modularizing such a network into individual modules by using the hybrid dynamic tree-cutting method. Modules were annotated by using three pathway databases, including Gene Ontology (GO), KEGG, and Hallmark, by Fisher’s exact test, and visualized by using Cytoscape.js.

### Ferrozine iron assay

Cells were seeded at 1 × 10^5^ per well of a 24-well plate and treated as described in the figure legend. Cells were treated with 100 μL 10 mM HCl and 100 μL of a solution of equal volumes of 1.4 M HCl and 4.5% potassium permanganate and incubated for 1 h at 60 °C within a fume hood. Then, 30 µL of detection reagent (6.5 mM ferrozine, 6.5 mM neocuproine, 2.5 M ammonium acetate, and 1 M ascorbic acid) was added to each tube. After a 30-min incubation, the solution was transferred into a well of a 96-well plate, and the absorbance was measured at 550 nm.

### Heme and porphyrin determination

Cells were pelleted and lysed in acetone acidified by 20% 1.6 N HCl for 5 min at room temperature and then centrifuged (21,000 × *g*) for 5 min at 4 °C. Then, 100 µL of supernatant was injected into the HPLC system (Shimadzu SIL20AC), and heme and PPIX were separated on a Supelco 125 × 4.6-mm C18 column (3 μm) by applying a 30–66% linear gradient mobile phase over 5 min followed by a 66– 90% linear gradient over 20 min. Heme absorbance was read at 400 nm wavelength, whereas porphyrin fluorescence was measured at wavelengths of 395 nm (excitation) and 630 nm (emission). Results were quantified by extrapolation to known quantities of external standards and normalized to cell number.

### Electron microscopy

Samples for electron microcopy were fixed as monolayers in a mixed aldehyde fixative in cacodylate buffer, then scraped and pelleted for further processing. Samples were post-fixed in osmium tetroxide and contrasted with uranyl acetate. Following dehydration in an ascending series of alcohols, samples were transitioned into EMbed812 resin with propylene oxide as the transitional solvent. Once in 100% resin, samples were polymerized overnight at 80°C. Samples were sectioned at ∼70-nm thickness on a Leica (Wetzlar, Germany) UC-7 ultramicrotome and examined in a ThermoFisher Scientific F20 transmission electron microscope. Images were captured on an AMT camera system (Woburn, MA). Unless otherwise stated, all reagents were sourced from Electron Microscopy Sciences (Hatfield, PA).

### Mitochondrial morphology

Images captured by electron microscopy (at least 16 representative fields with visible mitochondria) were analyzed for each treatment set of 3T3 cells: untreated or treated for 24 hours with ALA (200 µM), succinylacetone (250 µM) or with both. To avoid bias, samples were masked before scoring by the St. Jude Electron Microscopy Core. Mitochondria in each field (16-30 fields per treatment) were scored using the following scale: **0**-normal morphology mitochondria, **1**-no defect except a few cristae not extending across the mitochondria or minor variations in direction of cristae, **2**-many mitochondria with missing cristae, altered cristae direction, or highly unique/unusual mitochondrial shape 3-severely impaired mitochondria: most or all cristae missing or strongly altered, and/or mitochondria with membranes ruptured.

### Cell viability

Cells were seeded at either 2.5x10^3^ or 10^4^ cells per well of 96-well plates in 100-µL volume and treated with compounds as indicated in legends. Cell viability was determined by the use of an equal volume of Cell Titer Glo reagent (Promega) added directly to the media. After 5 minutes of mixing, the lysed cells in media/reagent solution were transferred to white plates (Thermo), and luminescence was determined in the Synergy H4 plate reader. Fluorescence was normalized to controls within each treatment group.

### Glutathione

Cells were seeded at 10^4^ per well of 96-well plates in 100-µL volume, and experimental wells were treated for 24 hours with 200 µM ALA alone or with other agents as indicated in legends. Total glutathione or GSSG was then measured by using the GSH/GSSH Glo kit (Promega) according to the manufacturer’s instructions. The fraction of oxidized glutathione was determined for each treatment group and normalized to untreated controls.

### Flow Cytometry

Cells were seeded at 2 x 10^5^ per well in 12-well plates and treated as indicated in legends for the times indicated. Cells remained in the dark with all fluorescent indicators followed by trypsinization and one wash with 1X PBS (Gibco). The indicators employed and fluorescence excitation/emission wavelengths were as follow: MitoSOX Red, mitochondrial superoxide indicator, 2 µM, (510/580 nm); DCFDA, general oxidative stress indicator, 5 µM, (485/535 nm); C11-Bodipy, lipid peroxidation indicator, 1 µM (488/545 nm); TMRE, mitochondrial potential indicator, 200 nM, (488/561 nm); Phen Green SK, cellular iron indicator, 10 µM, (400/510). Positive controls were added 30 minutes prior to staining and included antimycin A, 1µM, for MitoSOX Red; 0.5 mM hydrogen peroxide for DCFDA and C11-Bodipy; FCCP, 10 µM for TMRE; and 50 µM ferrous ammonium citrate for Phen Green SK. Background was subtracted using unstained cells, and mean fluorescence intensity of viable cells was normalized to stained controls, except for Phen Green. Because elevated iron concentration caused by Phen Green results in reduced fluorescence, these data are presented as inverse fluorescence.

### Cell surface labelling

Cells were seeded at 4 X 10^5^ density in 10 cm dish. When cells reached 90% confluency, they were treated with 200 µM ALA and 250 µM SA for 24 hours and then washed with PBX and harvested by trypsinization. The cells were then incubated with 0.25 mg/ml of EZ-Link Sulfo-NHS-SS-Biotin (Pierce) solution in PBS at 4°C for 30 minutes, with gentle mixing in a rotator. Excess labelling reagent was quenched by incubating the cells in 5mM Tris (pH 7.6). The cells were lysed in RIPA lysis buffer containing 1x protease inhibitor cocktail (Roche), 0.2 M phenylmethylsulfonyl fluoride (PMSF), 1M N-Ethylmaleimide (NEM) and 20 mM MG132. Protein quantity in the cell lysate was measured by BSA protein assay. Cell lysate containing 100 μg protein was incubated with 33 μl of streptavidin-agarose beads (Pierce) for 16-18 hours on a rotator at 4°C. The biotinylated surface proteins attached to the streptavidin agarose beads were eluted by boiling at 95°C for 5 minutes. The total eluates were analyzed by immunoblotting.

### Gene knockdown

NIH-3T3 cells were seeded in growth media at 2x10^5^ cells per well of a 6-well plate the day prior to siRNA transfection. Control and On-Target Plus siRNA pools directed against each target protein were obtained from Horizon/Dharmacon (Cambridge, UK), ferritin (L_047068_00_0020), Prdx3 (L_043352_01), and transferrin receptor (L_055550_01). Transfections for each well were performed with the desired siRNA in 150µl of media and combined with equal volume of OPTIMEM containing 9 µl Lipofectamine 2000. The final siRNA concentrations were 20 nM for ferritin and Prdx3 and 50 nM for the transferrin receptor. After 48 hours cells were split into 96-well plates as described in experiment legends or lysed for Western blotting. After attachment, cells were either left untreated or treated with a combination of 20 µM ferrostatin-1 and various concentrations of ALA. After an additional 48 hours viability was assessed using Cell Titer Glo reagent (Promega, Madison, WI) as per the manufacturer’s instructions.

### NMR Sample preparation and data acquisition and analysis of the Spectra

5 mg of the PPB was dissolved in 500 ml of DMSO-d_6_. All the NMR data were acquired on a Bruker NMR spectrometer operating at 700 MHz for proton resonance under Topspin (version 4.1, Bruker, Germany) with a triple resonance TCI cryoprobe at 298 K. For the proton assignment of the probe molecule, PPB, two-dimensional (2D) [^1^H, ^1^H] COSY, TOCSY, NOESY (mixing time of 70 msec) experiments were performed. The carbon assignments were confirmed with a 2D [^1^H, ^13^C] HSQC and a 2D [^1^H, ^13^C] HMBC experiment. The nitrogen chemical shifts were confirmed using a 2D [^1^H, ^15^N] HSQC. All the spectra were processed using nmrPipe and analyzed using Sparky. Since the product was dissolved in DMSO-d_6_ the exchange of the HN protons are fast and hence several expected NOEs could not be observed in the spectrum, irrespective of using long mixing times. Though the PPB had decayed over a period of storage (few unassigned peaks in the spectra), the assigned chemical shifts of all the protons (numbering as marked in Supplemental Data Figure 1) are given in Supplemental Data Table 1. Some of the quaternary carbon chemical shifts were derived from HMBC spectrum. Since the product was dissolved in DMSO-d_6_ the exchange of the HN protons are fast and hence several expected NOEs could not be observed in the spectrum, irrespective of using long mixing times. The NOE’s observed from H32 (3.63 ppm, methyl) to HN7(6.97 ppm) is an indicator of the PPIX linked to PEG in proximity, and the NOE’s from HN8 (6.95 ppm) to H81 (1.30 ppm), H49 (2.25 ppm) and H56 (0.765 ppm) show the connectivity between PEG and the Biotin moiety (Supplemental Data Figure 2).

## Funding Sources

This work was supported by NIH grants R01 CA194057, CA194206 (JS), P30 CA21745, CA21865, and CA96832, and by ALSAC. The Cellular Imaging and Flow Cytometry Shared Resources are supported by SJCRH and NCI P30 CA021765.

## Author Contributions

Conception and design: John D Schuetz and John Lynch Development and methodology: John Lynch, Junmin Peng, Yu Fukuda, Yao Wang, Sabina Ranjit, Camenzind G Robinson, Christy R. Grace, Youlin Xia, Kanisha Kavdia Analysis and Interpretation of Data: John Schuetz, John Lynch, Yuxin Xi, Christy R. Grace, Youlin Xia Writing and revision of manuscript: John D. Schuetz, John Lynch Data Availability: Proteomic data will be available through the Dryad data depository via doi:10.5061/dryad.mkkwh712t.

## Competing interests

John D. Schuetz on behalf of the authors declares that there are no competing financial interests in the development of this work.

## Supplemental Data

**Supplemental Fig. 1.**
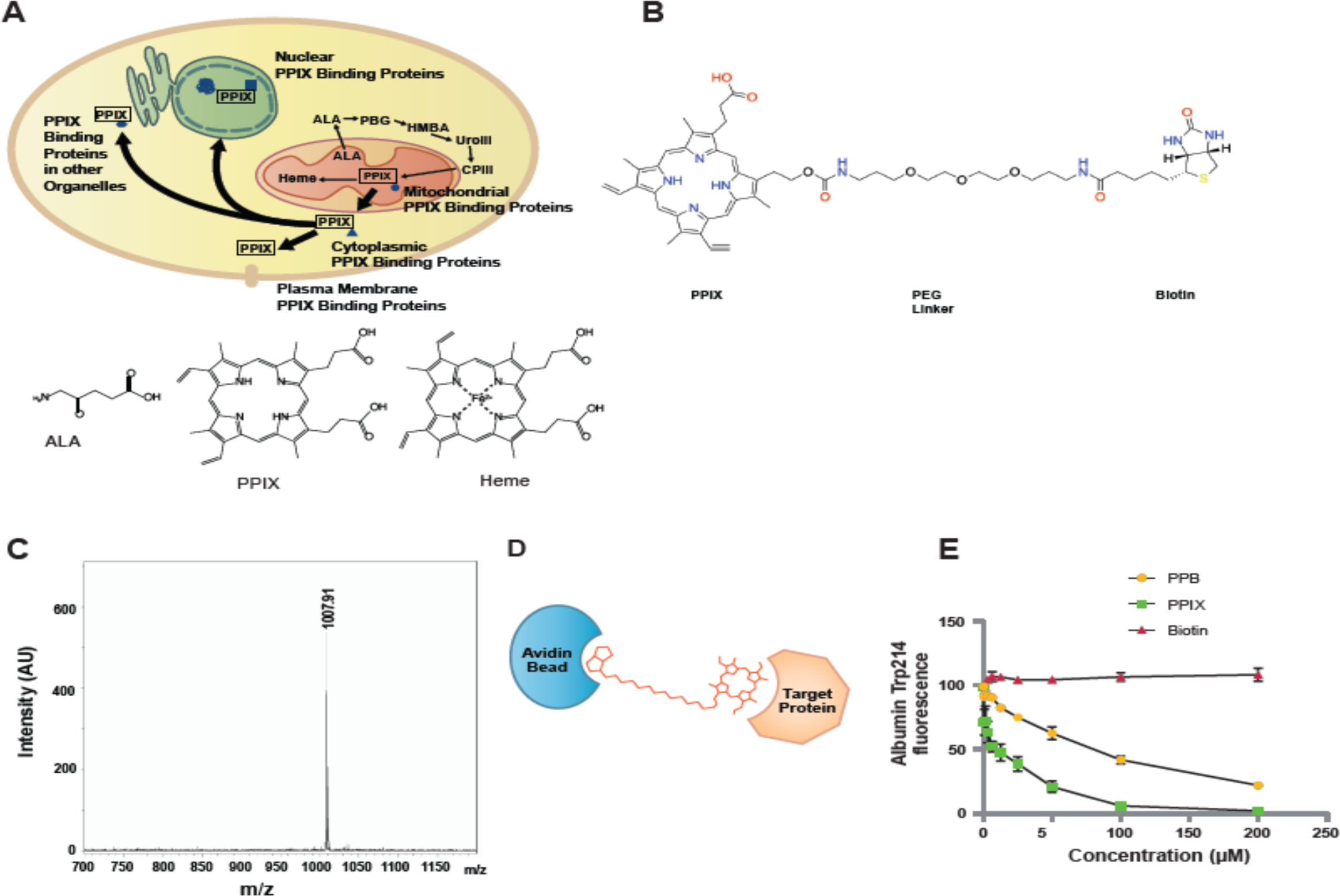
PPIX-binding protein method. **A,** PPIX is the penultimate intermediate in the heme biosynthetic pathway. It egresses from the mitochondria and is chaperoned to transporters to exit the cell. **B,** Diagram of the PPIX-binding (PPB) probe, which for simplicity, does not show all the PEG moieties. **C,** Mass spectrogram of the PPB probe showing a single molecular species. **D,** Depiction of the interaction between PPB, avidin-bead, and target protein. **E,** PPB was tested for binding with and quenching of the fluorescence of albumin (mean ± SEM), which is comparable to PPIX. (n=3 independent samples)

**Supplemental data Fig 2.**
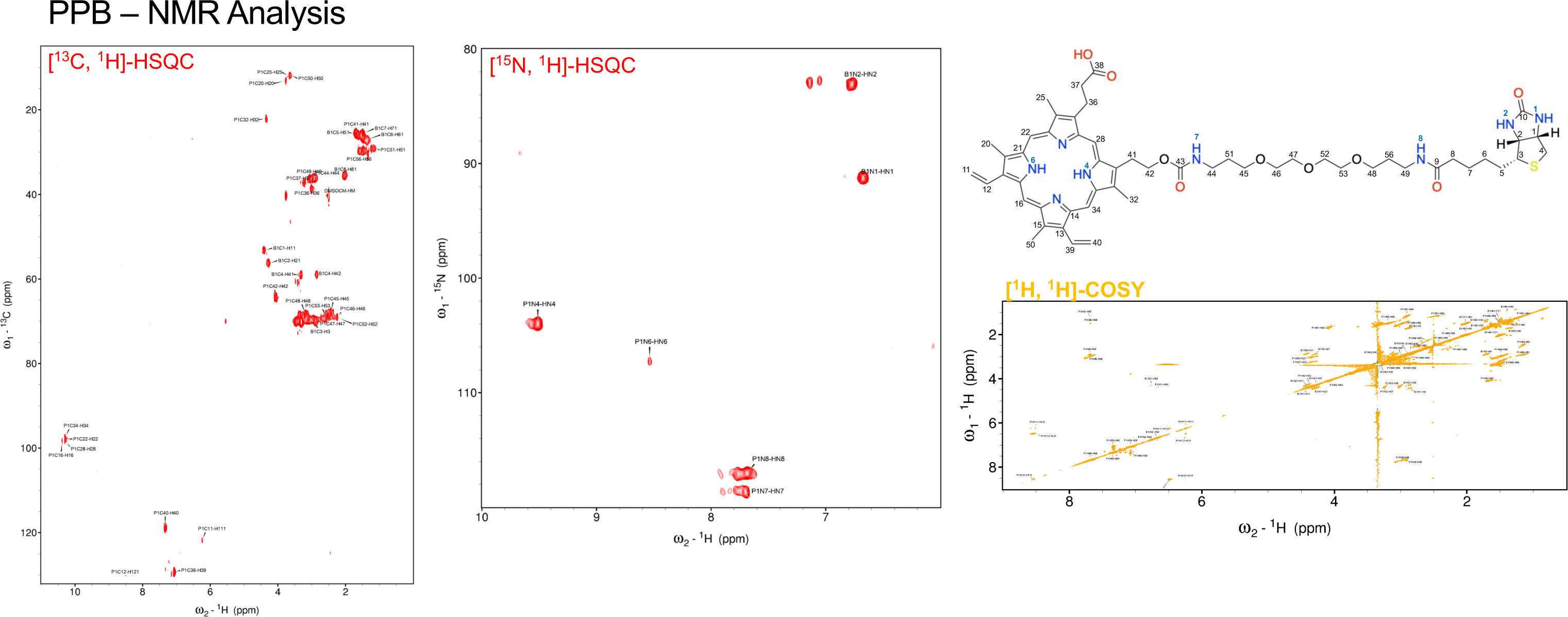
NMR analysis of PPB probe

**Supplemental data Fig 3.**
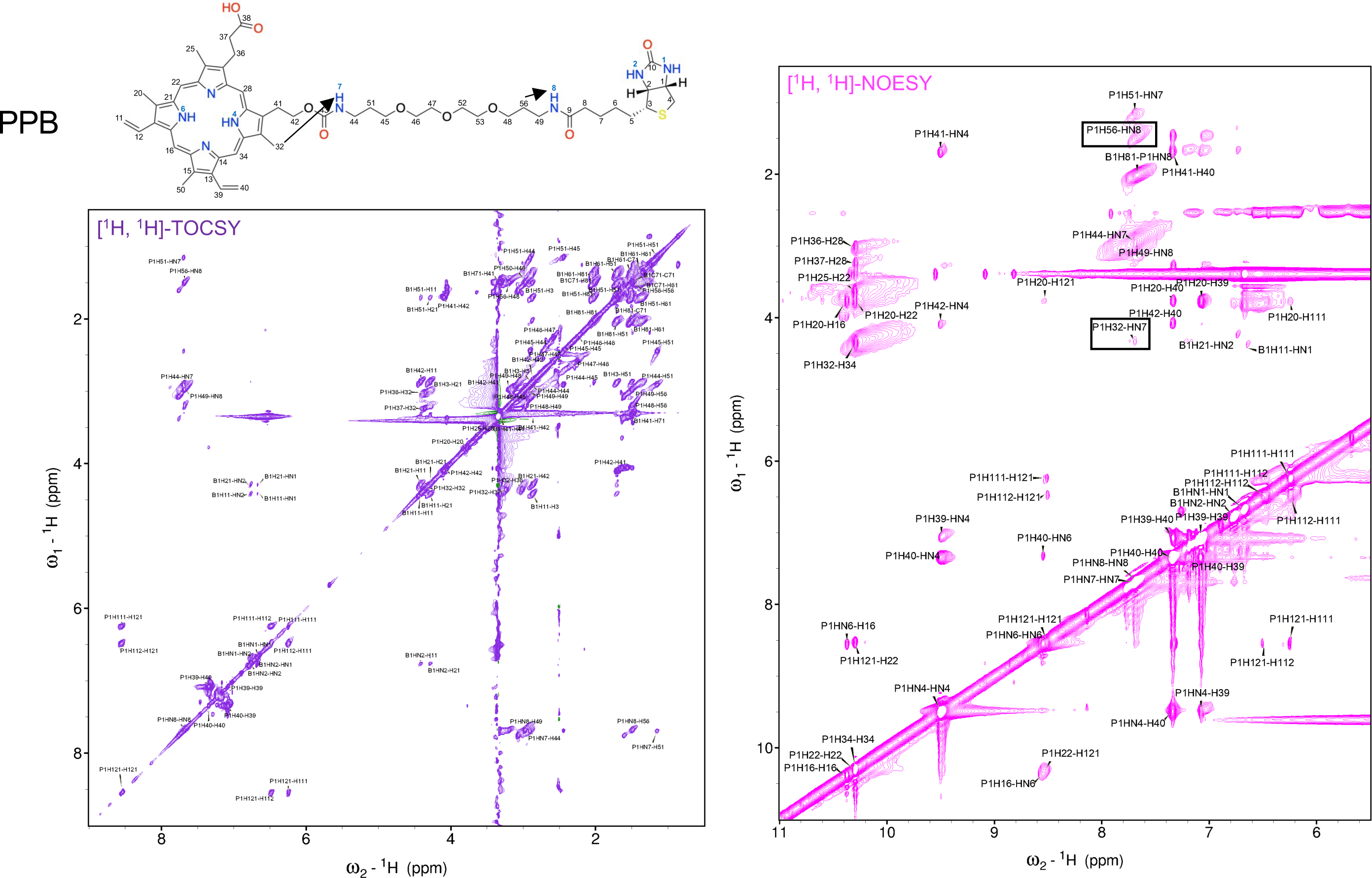
NMR analysis of PPB probe

**Supplemental Figure 4.**
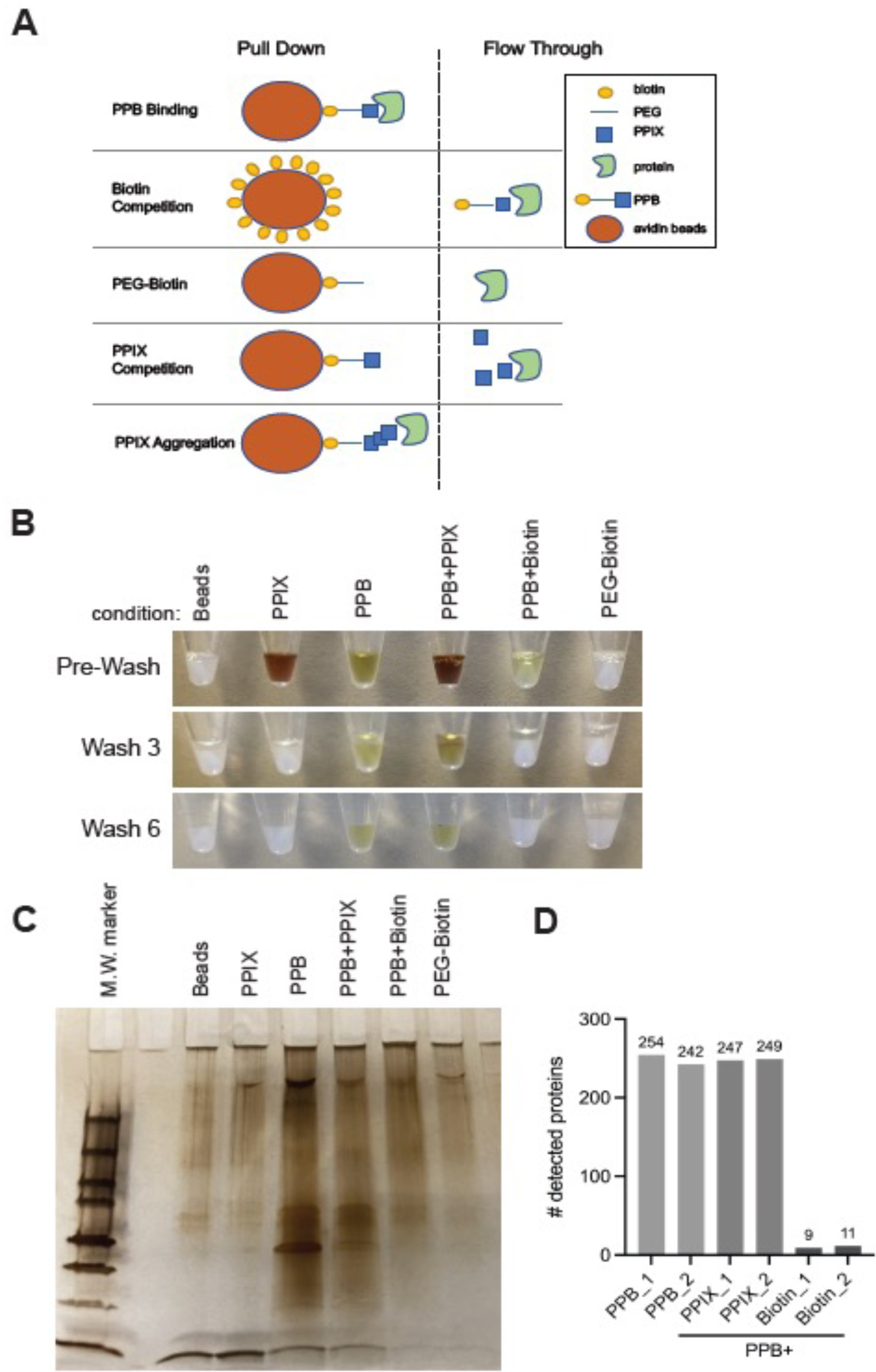
A). Model of predicted and observed interactions for PPB pulldown and in combination with competitive inhibitors. Streptavidin beads (orange circle) binds to the biotin moiety (yellow circle) of PPB. The heme moiety of PPB (blue square) binds to interacting proteins in the lysate (green polygon). B). Streptavidin bead coloration at various stages of washing for 0.5 mg cell lysate incubated with untreated beads or in combination with PPIX, PPB, PPIX/PPB coincubation, Biotin/PPB coincubation, or with PEG-Biotin. Color illustrates wash-resistant binding of PPIX to beads only in combination with PPB C). Silver stain of corresponding protein pulldowns from the above beads. D). Number of independent proteins detected by mass spectrometry in pulldowns with PPB alone versus with competition by PPIX (7.5X excess cotreatment) or biotin (50X pretreatment).

**Supplemental Data Fig. 5.**
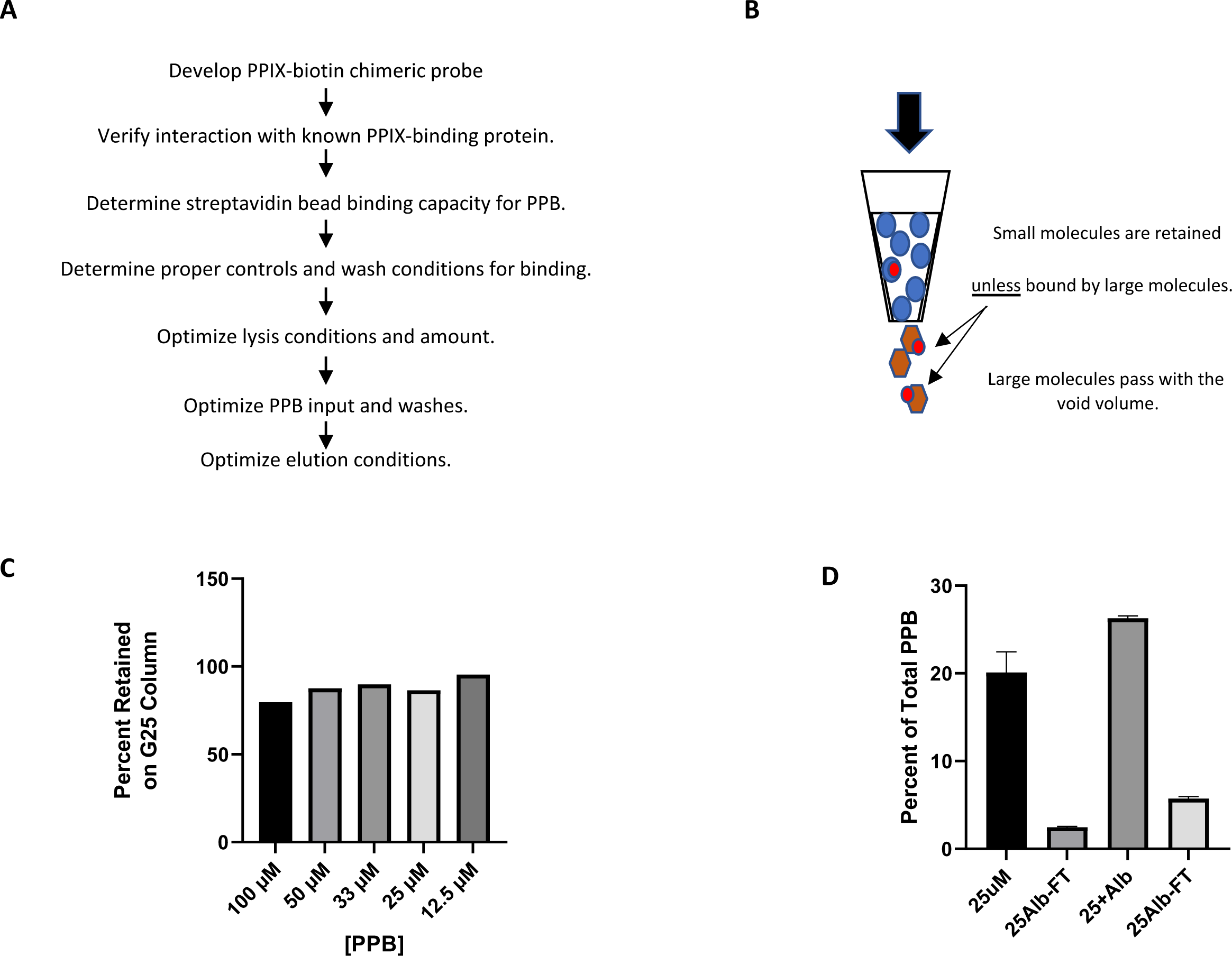
Design and testing of the PPIX-biotin (PPB) probe. **A**) Workflow for the development of PPIX-biotin probe and assay for subsequent pulldown experiments and analysis. **B**) Principle for testing albumin binding by PPB: albumin-bound PPB will pass through size-exclusion gel while free probe will be retained. **C**) PPB from 12.5-100 µM is retained by G25 column. **D**) Protein bound PPB passes through the column.

**Supplemental data Fig.6:**
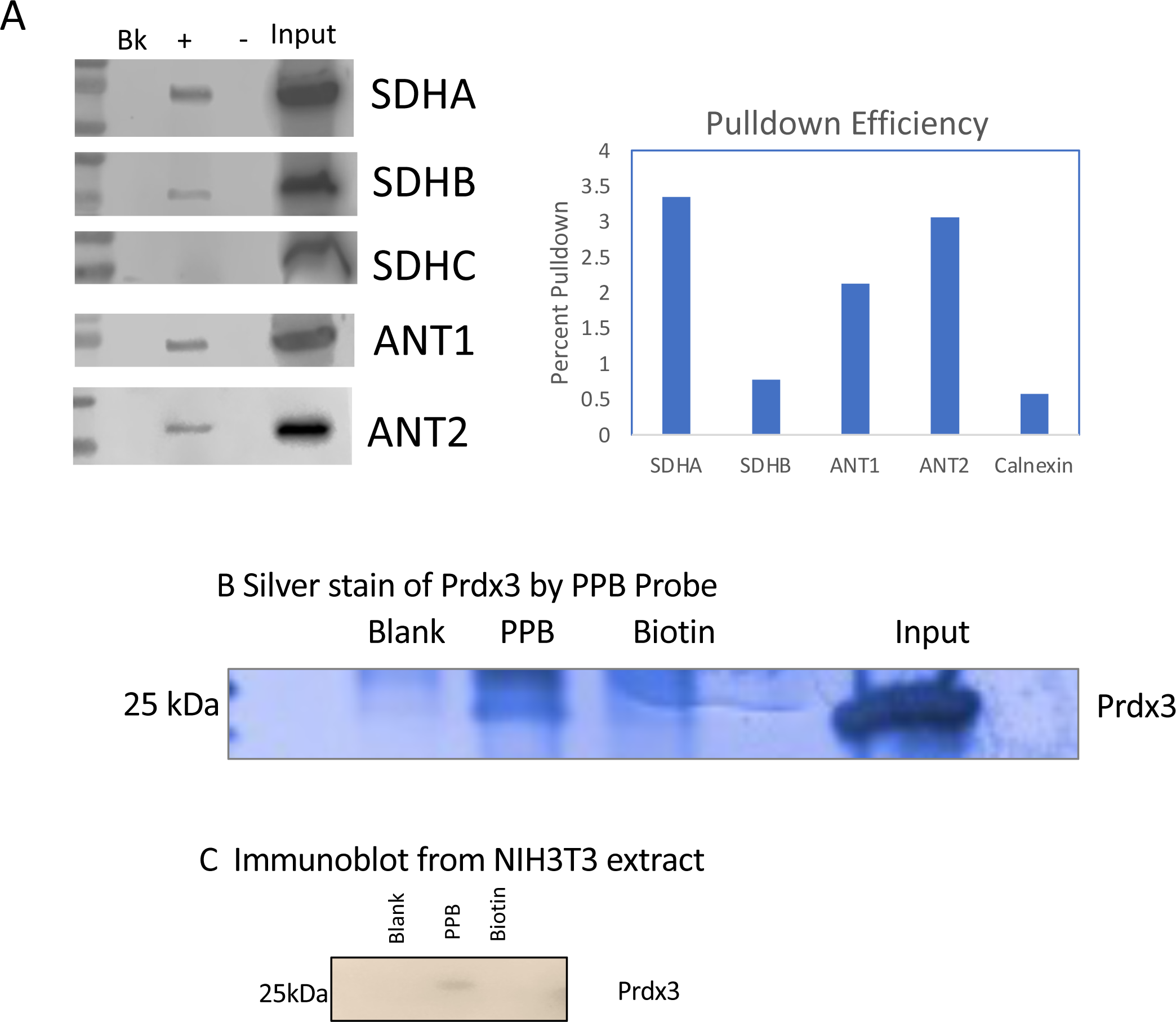
immunoblot confirmation of proteins Identified in PPB Proteomic Screen. **A)** 100 μg U251 lysate (input) was precleared 2X with washed streptavidin beads. The supernatant (post-clearance) was incubated with 20 nM PPB in the dark then bound with 100 μl of fresh beads. Supernatant (sup) was removed and beads washed 6X in PBS/0.1% TX100 and beads extracted in Laemmli buffer (pulldown) for 5 min at 95 C. The proteins were identified by immunoblot analysis and the efficiency of “pulldown” determined relative to input. **B)** Prdx3 was incubated with PPB and the binding competed with Biotin. The blank shows the residual protein trapped by the beads. **C**. Prdx3 is pulled down from an NIH3T3 extract.

**Supplemental Data Fig 7.**
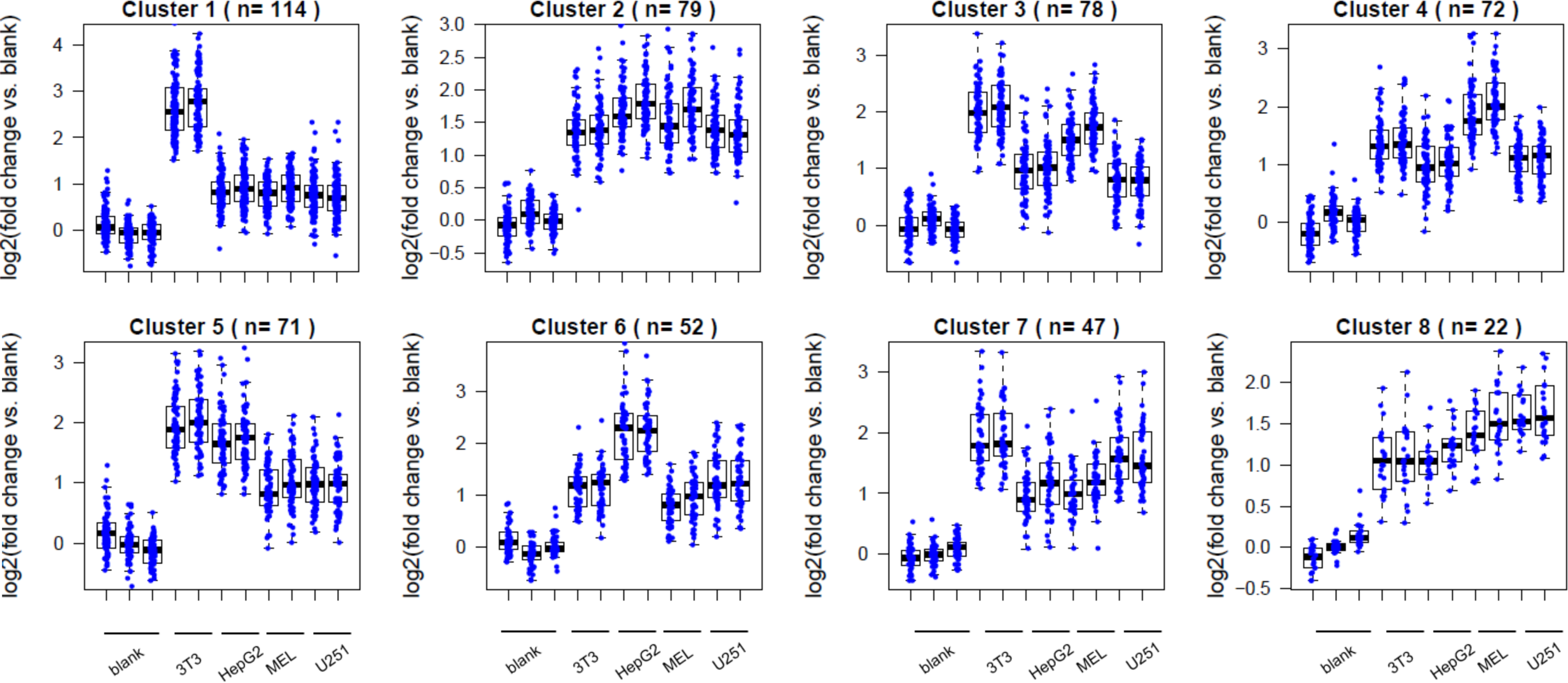
Relative level of protein clusters among cell lines. WGCNA (Weighted gene co-expression network analysis) of TMT quantitative proteomics data by PPIX pulldown identified eight distinct co-expression patterns across cell lines. Each dot represents one protein, with **the number of proteins within each cluster indicated in parentheses**.

**Supplemental Data Fig 8.**
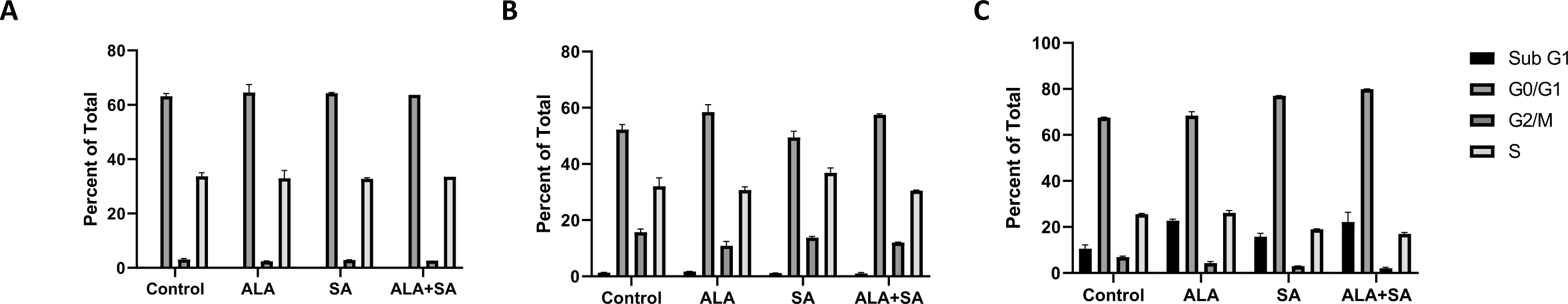
ALA treatment of NIH-3T3 cells does not alter cell cycle. NIH3T3 cells were treated with 200 µM ALA at A) 24 hours B) 48 hours or C) 72 hours progression, as determined by FACS by using propidium iodide.

**Supplemental Data Fig. 9.**
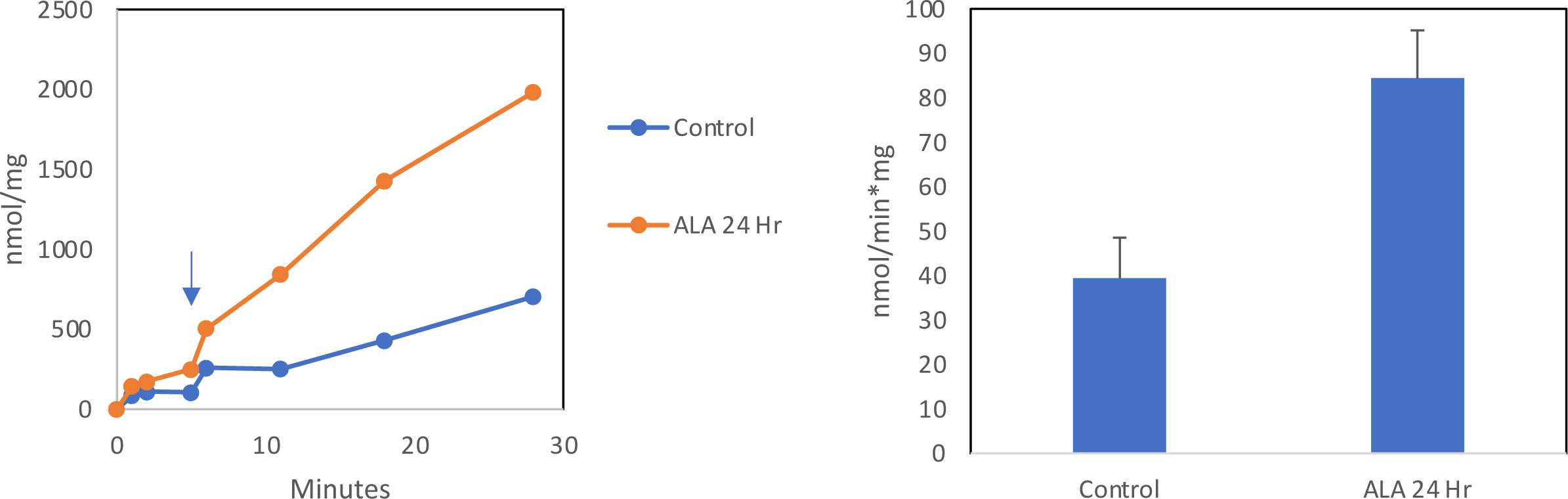
GPX4 Activity increases in NIH3T3 treated with ALA. ALA treatment 200 µM for 24 hours. GPX4 assay in 1 mM GSH, NaN3, EDTA, 0.2 mM NADPH, 3 units/ml glutathione reductase, 1mM cumene hydroperoxide.

**Supplemental Data Fig 10.**
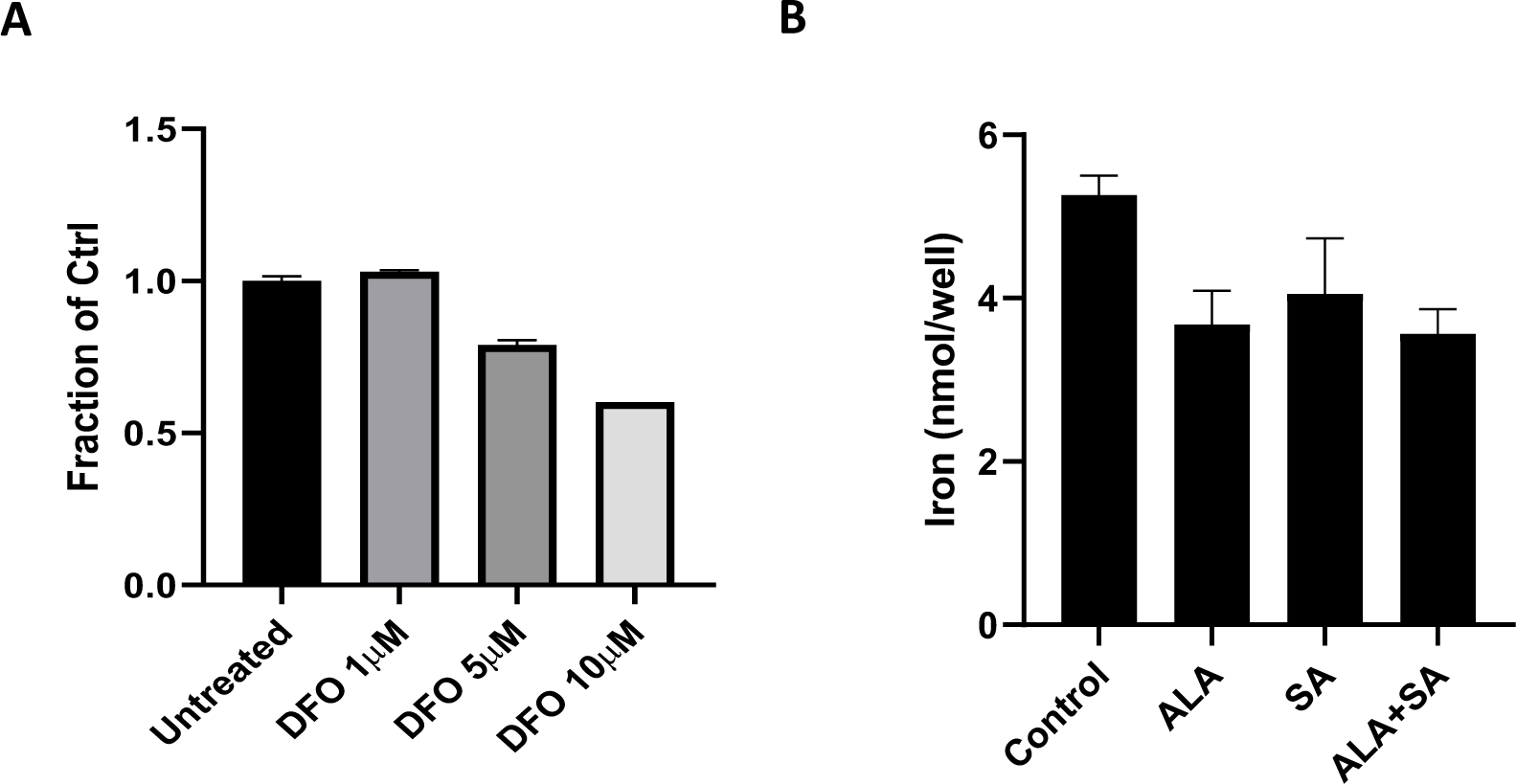
DFO chelation and intracellular iron. **A)** Desferrioxamine shows low levels of toxicity compared to untreated control NIH3T3 cells after 72 hr exposure. **B)** Free intracellular iron levels in NIH-3T3 cells are decreased in both ALA and ALA+SA treatments. Cells treated with ALA and/or succinylactone for 48 hours, with 40 µM FAC present for the final 24 hours, showed slightly reduced iron in treated wells, as determined by ferrozine assay.

**Supplemental Data Fig 11.**
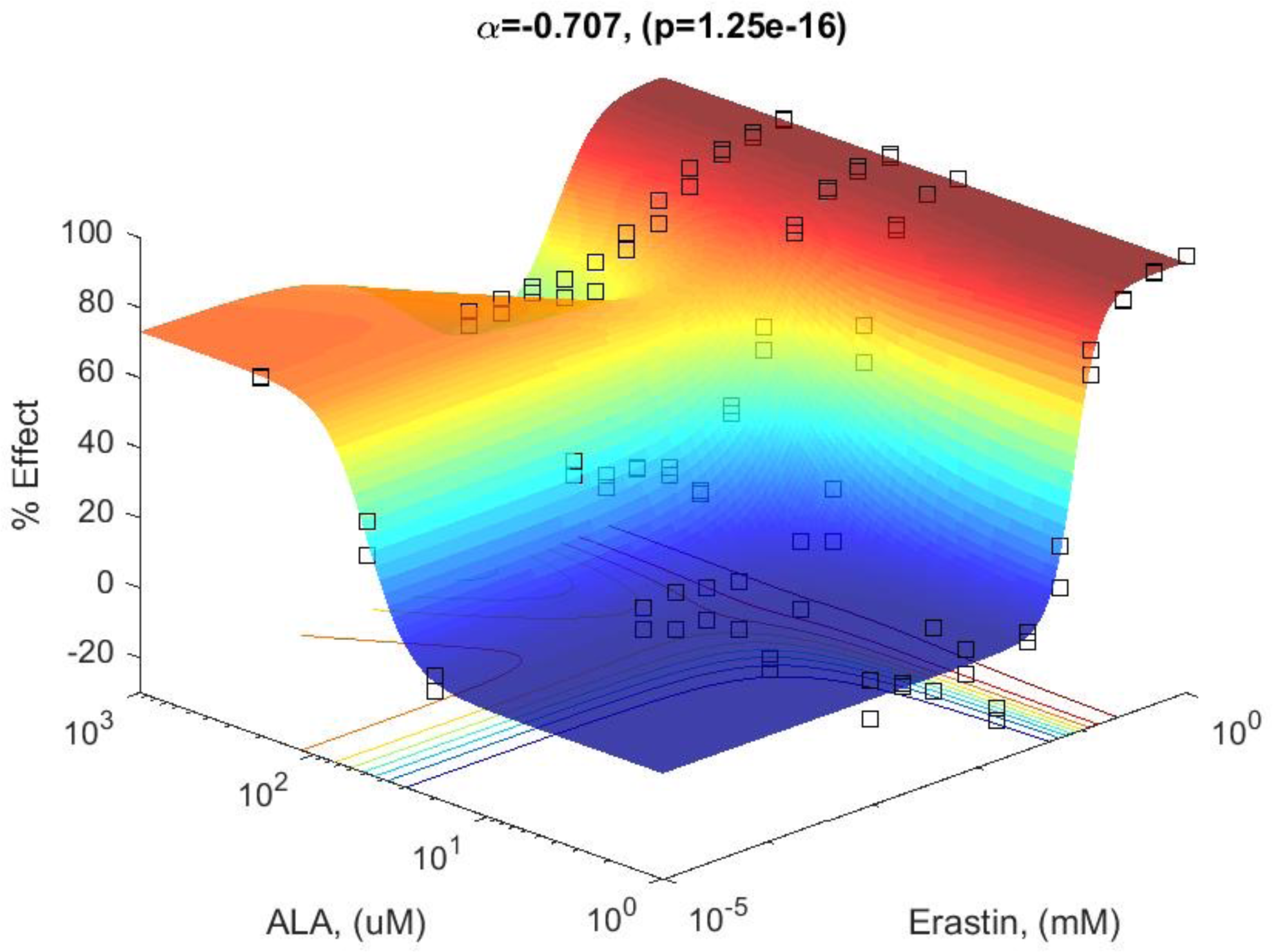
Analysis of synergy between ALA & Erastin.

**Supplemental Data Fig 12.**
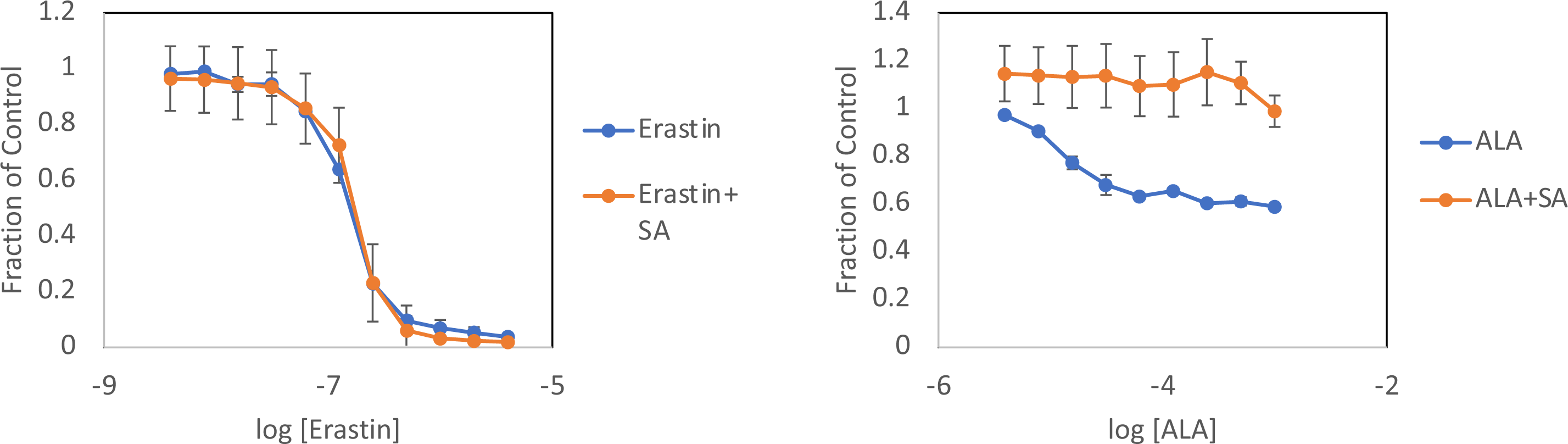
Succinylacetone rescues ALA, but not erastin, toxicity. Erastin toxicit at 48 hours is not affected by succyiacetone treatment a), however succinylacetone cotreatment produces near complete attenuation of ALA induced toxicity at the same treatment time.

**Supplemental Data Fig. 13.**
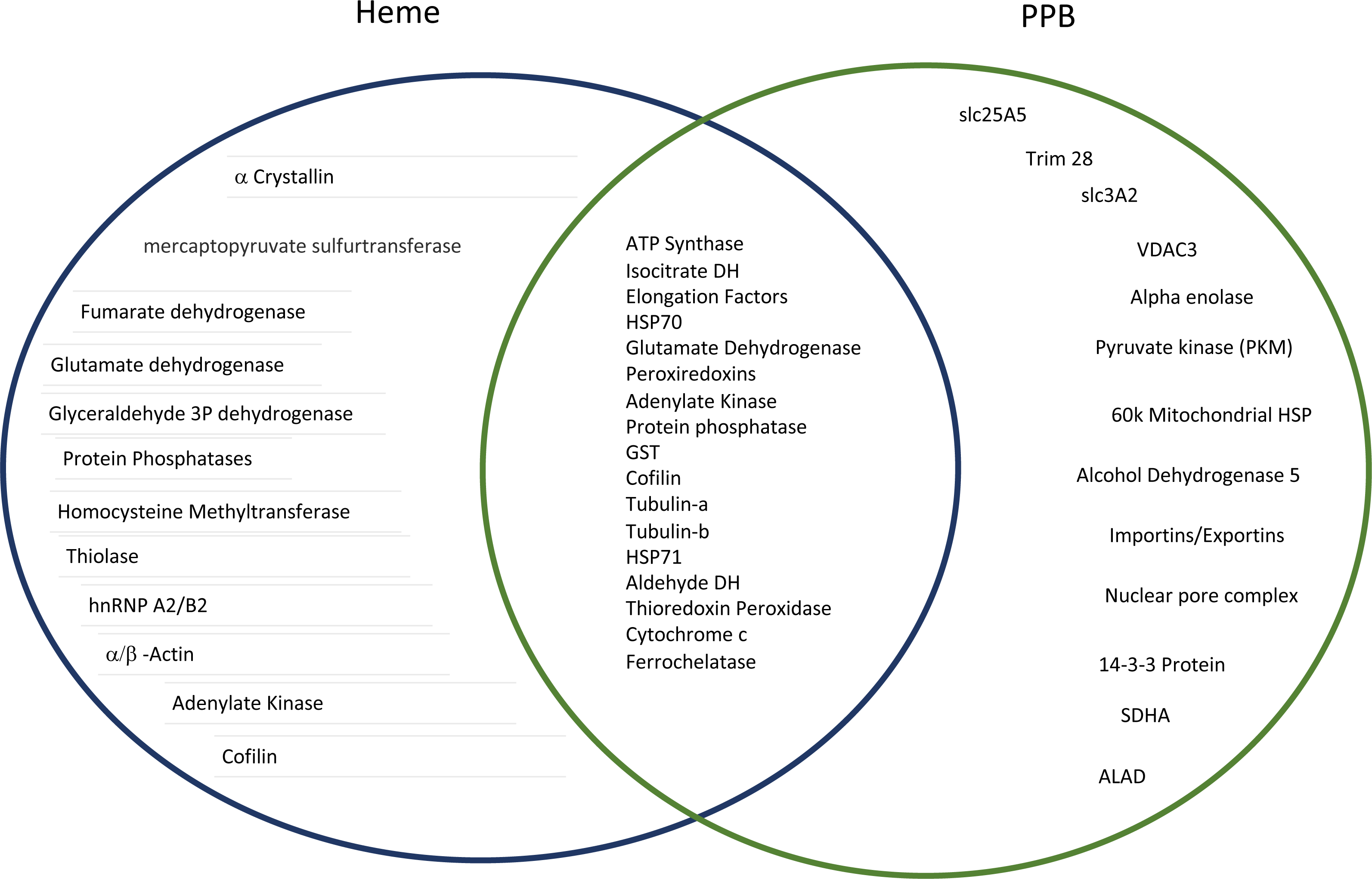
Heme and PPIX binding proteins by Venn diagram.

**Supplemental Data Table 1.** PPB chemical shifts at 298K

| Group | Atom | Shift | SDev | Group | Atom | Shift | SDev |
| --- | --- | --- | --- | --- | --- | --- | --- |
| B1 | C1 | 53.171 | 0.015 | P1 | C45 | 67.706 | 0.045 |
| B1 | C10 | 161.686 | 0.022 | P1 | C46 | 68.876 | 0.01 |
| B1 | C2 | 56.24 | 0.044 | P1 | C47 | 69.363 | 0.026 |
| B1 | C3 | 70.156 | 0 | P1 | C48 | 68.385 | 0.021 |
| B1 | C4 | 59.067 | 0.008 | P1 | C49 | 36.126 | 0.033 |
| B1 | C5 | 25.648 | 0.02 | P1 | C50 | 11.868 | 0 |
| B1 | C6 | 27.089 | 0 | P1 | C51 | 29.123 | 0.026 |
| B1 | C7 | 25.591 | 0.019 | P1 | C52 | 69.122 | 0 |
| B1 | C71 | 1.504 | 0.002 | P1 | C53 | 69.123 | 0 |
| B1 | C8 | 35.524 | 0 | P1 | C56 | 29.73 | 0.007 |
| B1 | H11 | 4.418 | 0.003 | P1 | C9 | 172.384 | 0.015 |
| B1 | H21 | 4.286 | 0.005 | P1 | H111 | 6.244 | 0.003 |
| B1 | H3 | 2.879 | 0.005 | P1 | H112 | 6.481 | 0.006 |
| B1 | H41 | 3.317 | 0.002 | P1 | H121 | 8.53 | 0.018 |
| B1 | H42 | 2.862 | 0.003 | P1 | H16 | 10.381 | 0.005 |
| B1 | H51 | 1.709 | 0.002 | P1 | H20 | 3.77 | 0.006 |
| B1 | H61 | 1.398 | 0.004 | P1 | H22 | 10.302 | 0.006 |
| B1 | H71 | 1.493 | 0.003 | P1 | H25 | 3.64 | 0.008 |
| B1 | H81 | 2.023 | 0.003 | P1 | H28 | 10.292 | 0.003 |
| B1 | HN1 | 6.672 | 0.001 | P1 | H32 | 4.353 | 0.005 |
| B1 | HN2 | 6.766 | 0.003 | P1 | H34 | 10.289 | 0.006 |
| B1 | N1 | 91.286 | 0 | P1 | H36 | 3.016 | 0.003 |
| B1 | N2 | 83.093 | 0 | P1 | H37 | 3.24 | 0.003 |
| P1 | C11 | 121.832 | 0 | P1 | H39 | 7.079 | 0.006 |
| P1 | C12 | 130.696 | 0 | P1 | H40 | 7.341 | 0.004 |
| P1 | C13 | 135.62 | 0 | P1 | H41 | 1.693 | 0.004 |
| P1 | C14 | 143.86 | 0 | P1 | H42 | 4.094 | 0.005 |
| P1 | C15 | 137.565 | 0 | P1 | H44 | 2.909 | 0.002 |
| P1 | C16 | 98.266 | 0 | P1 | H45 | 2.442 | 0.003 |
| P1 | C19 | 129.292 | 0 | P1 | H46 | 2.254 | 0.003 |
| P1 | C20 | 13.231 | 0 | P1 | H47 | 2.586 | 0.004 |
| P1 | C21 | 135.929 | 0 | P1 | H48 | 3.193 | 0.003 |
| P1 | C22 | 97.866 | 0 | P1 | H49 | 2.972 | 0.003 |
| P1 | C25 | 11.858 | 0 | P1 | H50 | 3.114 | 0.935 |
| P1 | C28 | 98.312 | 0 | P1 | H51 | 1.149 | 0.003 |
| P1 | C32 | 22.119 | 0.011 | P1 | H52 | 2.249 | 0 |
| P1 | C34 | 97.448 | 0 | P1 | H53 | 2.573 | 0 |
| P1 | C36 | 38.699 | 0 | P1 | H56 | 1.492 | 0.002 |
| P1 | C37 | 37.184 | 0 | P1 | HN4 | 9.499 | 0.006 |
| P1 | C38 | 174.647 | 0 | P1 | HN6 | 8.546 | 0.005 |
| P1 | C39 | 129.379 | 0.09 | P1 | HN7 | 7.692 | 0.002 |
| P1 | C40 | 118.779 | 0.004 | P1 | HN8 | 7.68 | 0.004 |
| P1 | C41 | 25.69 | 0.001 | P1 | N4 | 103.985 | 0 |
| P1 | C42 | 64.131 | 0.016 | P1 | N6 | 107.255 | 0 |
| P1 | C43 | 154.072 | 0 | P1 | N7 | 118.693 | 0 |
| P1 | C44 | 36.056 | 0.025 | P1 | N8 | 117.032 | 0 |

## Notes

### Competing Interest Statement

The authors have declared no competing interest.

